# SOPSim: A cell vertex-based application for simulating sensory bristle patterning with dynamic filopodia

**DOI:** 10.64898/2026.11.04.764402

**Authors:** William Ebo Annan, Olaoluwayimika Olugbenle, Ginger Hunter, Emmanuel Asante-Asamani

## Abstract

Repeating patterns, such as the bristles on the thorax of fruit flies developed from sensory organ precursor (SOP) cells, play important roles in the lives of flies as they are connected to the central nervous system and help detect environmental cues and threats. Notch signaling is known to regulate the formation of these patterns. Primarily, Notch signaling involves local communication between neighboring cells in direct contact (*apical or junctional interaction*). However, this short-range interaction alone cannot fully explain the complex biological patterns observed. Further studies have revealed long-range communication between cells through actin-based protrusions known as filopodia. The precise understanding of how the dynamics of filopodia—such as protrusion and retraction—lead to Notch–Delta activation remains unclear, which limits our ability to fully comprehend the mechanisms underlying cell fate decisions and the emergence of spatially organized patterns during tissue development and regeneration.

In this study, we extend the modeling framework for Notch signaling by incorporating a filopodia dynamics model driven by myosin-mediated contraction. We address existing gaps by defining criteria under which any given pair of filopodia will physically intersect. We also introduce the local sensory area (LSA), a metric for quantifying local SOP spacing, and use experimental data to calibrate a Notch threshold that distinguishes SOP from epithelial cells. Through numerical simulations, we confirm that long-range communication via filopodia is essential for reproducing the refined patterns observed *in vivo*. Additionally, we explore how filopodia length, lifetime, and orientation influence the spatial organization of SOP cells. To this end, a Python library has been developed which allows users to perform simulations of Notch Delta signaling via a (1) Graphical User Interface/website, local to the user’s computer, (2) via commands in the command line interface, or (3) by calling the simulation using a function via a Python API. This library allows users without computational expertise to easily specify a small number of input parameters and generate simulations of the model.

## 1 Introduction

Biological patterns such as stripes on zebra [1], leopard spots [2–4], bristle cells on the thorax of fruit flies [5–7] among many others play essential roles in the survival of organisms. For predators, these patterns aid in camouflage, allowing them to approach prey undetected, while for prey, the patterns help them evade detection. In fruit flies, bristle cells on the thorax are connected to the central nervous system and assist in sensing environmental cues to avoid threats [8, 9]. These patterns are formed during early stages of development [10–12]. In this study, we focus on the formation and organization of sensory organ precursor (SOP) cells, which develop to form the bristles on the thorax of fruit flies.

Bristle cell formation occurs through a contact-based signaling mechanism that involves interactions between cells during developmental stages, specifically during the transition from the pupal to adult stage [13–16]. Immediately after pupal formation, the cells within the tissue express a fairly uniform distribution of Delta ligands and Notch receptors [17, 18]. During cell-to-cell interaction, a Delta ligand from one cell, referred to as the *signal-sending* cell, binds to a Notch receptor on a neighboring cell, referred to as the *signal-receiving* cell. This binding can result in Notch activation, where the Notch receptor is cleaved at the plasma membrane of the signal-receiving cell, separating the Notch intracellular domain (NICD) from the Notch extracellular domain (NECD) [19–22]. Following the Notch activation, the NECD is endocytosed by the signal-sending cell [23–25], while the NICD is freed into the cytoplasm and ultimately enters the nucleus of the signal-receiving cell [25, 26]. Inside the nucleus, the NICD modulates specific gene activities, including repressing expression of Delta in the signal-receiving cell [19, 26–28], while promoting expression of Notch. As development progresses, some cells end up expressing more Delta, while others accumulate more Notch. Cells that express higher Delta become SOP cells, which later undergo cell division to form bristles, whiles cells that accumulate more Notch become epithelial cells [15, 29, 30]. Notch signaling occurs through direct contact between adjacent cells, known as *apical or junctional interaction*, and through long-range interactions along basal cellular protrusions called *filopodia*, which allows non-adjacent cells to communicate [14, 15, 31, 32]. While apical interaction is well understood, the mechanism behind long-range interaction via filopodia remains unclear [33–35]. For example, it is not yet fully understood how Notch activation occurs along filopodia At apical or junctional interactions, endocytosis of Delta helps to activate Notch; it is not currently known if Delta endocytosis occurs on filopodia.We also don’t know the mechanism by which NICD is transported along filopodia to the cell cytoplasm and the rate at which NICD enters the cell body at the base of filopodia. The influence of all these subprocesses on pattern formation remains to be studied. Understanding the formation and spatial organization of SOP cells is important for elucidating the mechanisms underlying cell fate determination and tissue patterning. In particular, it provides insight into how lateral inhibition, mediated through filopodia dynamics, regulates Notch–Delta signaling and contributes to the emergence of organized developmental patterns. Such understanding may also help clarify how genetic perturbations influence cellular behavior and tissue organization

Existing mathematical models of Notch–Delta signaling do not fully capture the protrusion and retraction dynamics of filopodia, which could be critical for studying the effect of long range activation and transport of Notch on SOP patterning. The pioneering mathematical model developed by J. Collier et al. [34] considered only apical interactions between neighboring cells. Later, M. Cohen et al. [15] extended this framework by incorporating long-range basal interactions mediated by filopodia and demonstrated that such interactions are necessary for reproducing the refined patterns observed in wild-type tissues. Although their model represented an important advancement, the underlying dynamics of filopodia were treated in a simplified, semi-heuristic manner, without the ability to specify precise length dynamics over typical protrusion lifetimes. Experimental studies have established that filopodia continuously undergo cycles of protrusion, sometimes pausing and retraction throughout their lifetime [36–45]. Furthermore, microscopic observations show that even cells in direct contact can interact through basal filopodia. These behaviors were not explicitly represented in the simulations of Cohen et al. [15]. In their model, filopodia turnover was introduced probabilistically: at each time step, a random number was generated and compared against a prescribed rate to determine whether a filopodium expired.

Expired filopodia were then replaced with new filopodia whose lengths were randomly sampled from a normal distribution with mean *F*_*µ*_ and standard deviation *F*_*σ*_. While this approach captures stochastic turnover, it does not describe the continuous length dynamics arising from protrusion, sometimes pausing and retraction processes, limiting the use of this model to study how long range Notch activation can affect patterning. In addition, the two-dimensional simulations in [15] assumed that cells in the first surrounding layer interact exclusively through apical contact, whereas cells in the other layers interact only through filopodia. This assumption overlooks the possibility that cells in direct contact may also communicate through basal filopodia interactions and therefore limits the biological realism of the model.

Experimental evidence also indicates that filopodia exhibit heterogeneous lengths and lifetimes [46, 47]. Despite this, the precise role of filopodia length and lifetime in regulating spatial organization has not been completely characterized. Notch activation in a given cell depends on the amount of Delta expressed by neighboring cells. To account for the fact that not all interactions lead to signaling activation, Cohen et al. [15] introduced probability weights *α*_*a*_ and *α*_*b*_ to scale the contributions from apical and basal neighbors, respectively. However, the biological basis for the choice of these weights remains unclear. Similarly, different studies adopt different threshold values of Notch and Delta expression to identify cell state transition to epithelial or SOP fate [15, 48, 49], with limited justification for the selected thresholds. In addition, there is limited quantitative characterization of tissue-wide SOP spacing for the two-dimensional tissue. In two-dimensional tissues, SOP organization is commonly quantified using the fraction of SOP cells in the tissue referred to as the **relative sensory area** (RSA) [15]. Although this measures the abundance of SOP cells, it does not capture their spatial arrangement. For example, two tissues with the same RSA may exhibit fundamentally different spatial organizations, such as clustered versus uniformly distributed SOP cells. Proper tissue-wide quantification of SOP organization could reveal more intricate effects of Notch-signaling on patterning.

In this study, we extend the Notch–Delta signaling framework developed by Cohen et al. [15] by incorporating a model for filopodia length dynamics based on protrusion and retraction mechanisms. We also establish a deterministic criterion that governs whether two filopodia intersect within the tissue. By integrating these components into the existing signaling framework, the model more accurately represents the dynamic behavior of basal filopodia and provides a closer approximation of *in vivo* interactions. Using this extended framework, we investigate how filopodia length, lifetime and orientation influence the spatial organization of SOP cells. Furthermore, we introduce a criterion for cell fate determination informed by experimental Notch expression data from fruit flies. By varying the probability weights associated with apical and basal signaling interactions and comparing the resulting RSA values with experimental observations, we identify parameter ranges that produce spatial patterns consistent with wild-type tissues. To better quantify spatial organization, we introduce a new metric referred to as the *Local Sensory Area* (LSA). Unlike RSA, which measures only the global fraction of SOP cells, the LSA characterizes the local spatial distribution of SOP cells within the tissue. For each SOP cell, the LSA measures how neighboring SOP cells are distributed around it, thereby providing additional information about clustering and spatial uniformity. To make the result of this study more accessible to users without computational expertise, a Python library has been developed to perform Notch Delta signaling via a (1) Graphical User Interface (GUI)/website, local to the user’s computer, (2) via commands in the command line interface, or (3) by calling the simulation using a function via a Python API where users can specify a few parameters and generate simulations of the model

The remainder of this paper is organized as follows. In Section 2, we present the Notch–Delta signaling model and the model for filopodia length dynamics, modified from the earlier work of Mogilner et al. [50]. The filopodia model incorporates protrusion and retraction driven by retrograde flow and myosin-mediated contraction. We outline the detailed steps involved in implementing the coupled filopodia and Notch–Delta signaling models, including tissue creation, identification of neighbors and filopodia vectors, the deterministic condition governing filopodia intersection, and the definition of the *Local Sensory Area* (LSA), an alternative measure of SOP spacing. In Section 3, we present the results and discussion, examining the effects of the probability weights *α*_*a*_ and *α*_*b*_ on SOP spacing and investigating how filopodia length and lifetime influence pattern formation. Finally, conclusions and directions for future research are presented in Section 4.

## 2 Design and Implementation

Here, we describe the Notch–Delta signaling and filopodia dynamics models. We then describe the detailed implementation of the model, including tissue creation, identification of neighboring cells and filopodia intersections, computation of the effective Delta, and simulation of the Notch–Delta interaction model. We also describe the quantification of SOP spacing and the python library with graphical user interface (GUI) SOPSim which enables non-expert users to run simulations by specifying few model parameters.

### 2.1 Notch mediated lateral inhibition model

We adopt the model for Notch mediated lateral inhibition first introduced by Collier et al. [34] in 1996, and subsequently modified by Cohen et al. [15] in 2010 to incorporate the effects of filopodia. The modified model is given by

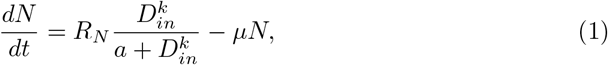

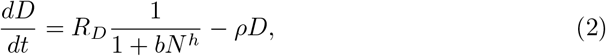

where

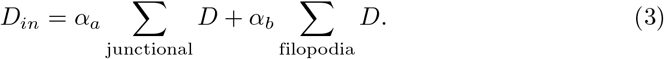

Here, *R*_*N*_ and *R*_*D*_ represent the production rate constants of Notch and Delta, respectively, while *µ* and *ρ* denote their corresponding decay rates. The term *D*_*in*_ represents the total effective Delta received from neighboring cells. In this framework, the incoming delta concentration *D*_*in*_ promotes the production of intracellular Notch *N* through Notch receptor activation. This is captured through a saturation term with Hill coefficient *k*. As Notch is produced, the intracellular concentration of Delta *D* is inhibited. The constants *h, a*, and *b* are parameters governing the changes in Notch and Delta concentrations within the cell. Here, Notch activation is not modeled explicitly, however the total Delta responsible for Notch activation *D*_*in*_ is taken to be proportional to the total intercellular Delta from adjacent cells (junctional), with an activation likelihood parameter *α*_*a*_, as well as the the intercellular Delta from cells reachable through filopodia interactions, with activation likelihood *α*_*b*_. Consequently, the effective Delta responsible for Notch activation in a given cell depends, in part, on the filopodia dynamics of neighboring cells. In Section 2.2, we present a deterministic model governing filopodia dynamics, to replace the semi-heuristic framework used by Cohen et al.

### 2.2 Filopodia dynamics model

Filopodia undergo a cycle of protrusion (extension), occasional pausing, and retraction. filopodia protrusion is primarily driven by the polymerization of actin monomers (G-actin) into filamentous actin (F-actin) at the leading edge of the protrusion. This assembly process generates forces that push the membrane outward, forming thin finger-like projections known as filopodia. The process is regulated by actin-nucleating proteins such as formins and Ena/VASP, which promote unbranched filament elongation [36–38].

Retraction occurs primarily through actomyosin contractility, in which myosin motors generate pulling forces that shorten and collapse the actin filament bundle within the filopodium [51]. This process is further facilitated by actin filament depolymerization and turnover, which disassemble F-actin into G-actin monomers that can be recycled for future filament growth [42–45, 52, 53]. The change in filopodia length *L* is therefore governed by the difference between the protrusion and retraction rates and can be represented mathematically as

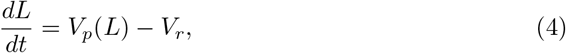

where *V*_*p*_(*L*) denotes the protrusion (extension) rate, and *V*_*r*_ denotes the retraction (retrograde flow) rate.

In 2005, Mogilner et al. [50] developed a model to study actin bundle mechanics and computed the minimum number of bundled actin filaments required to generate sufficient force for a filopodium to protrude against membrane resistance. Their model for the protrusion rate *V*_*p*_ is

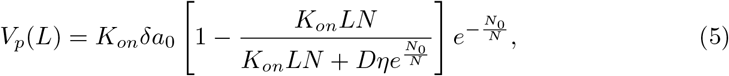

where, *K*_*on*_ is the assembly rate of G-actin, *δ* represents half the size of an actin monomer, *N* is the number of filaments in the filopodium, *D* is the effective G-actin diffusion coefficient, and *η* is the geometric conversion coefficient. The parameter

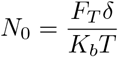

represents the minimum number of filaments required to support the filopodium, where *F*_*T*_ is the membrane tension and *K*_*b*_*T* is the thermal energy. The quantity *a*_0_ denotes the concentration of actin monomers at the base of the filopodium. We modified the G-actin assembly rate *K*_*on*_ to be non-constant, allowing for a reduced assembly during the retraction phase of the filopodia life cycle. We modeled *K*_*on*_ as follows

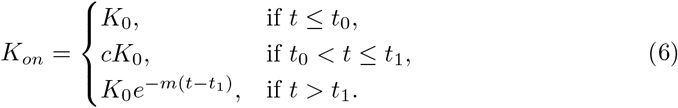

where *K*_0_, *m, t*_0_, *t*_1_, and *c ∈* (0, 1) are constants. Here, the assembly rate is assumed to be constant during protrusion, scales by some fraction *c* during a stationary phase and is reduced exponentially during retraction. The retraction rate of filopodia depends on the myosin pulling force (*F*_*m*_) [54–56], the membrane tension (*F*_*T*_) [42–45], and the drag coefficient of the filopodium (*τ*). Assuming that the filopodia behaves like a viscous fluid during retraction, we can model the retrograde flow velocity as

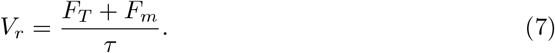

Since the dynamics of filopodia in the tissue can be expected to vary, we introduce variability in the length distribution of filopodia, defining

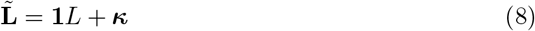

where **1** is an *N*-dimensional vector of ones, with *N* representing the number of filopodia in the tissue. The vector ***κ*** = (*κ*_1_, …, *κ*_*N*_)^*T*^ is defined component-wise by

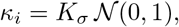

so that each *κ*_*i*_ is an independent sample from a standard normal distribution scaled by *K*_*σ*_.

### 2.3 Detailed Implementation

The implementation of the Notch–Delta signaling model involves tissue creation, identification of neighboring cells, computation of the effective Delta, and simulation of the filopodia dynamics model, which together are used to solve the Notch–Delta interaction model presented in Equation (1) and Equation (2).

#### 2.3.1 Tissue Creation

To construct the tissue, we assume that each cell is hexagonal with circumradius *r*, as illustrated in Figure 1B. The tissue is square with *N*_*c*_ cells in a row, as shown in Figure 1A. The centers of the cells in the first row are assumed to lie on the *x*-axis. The vertical distance between the centers of two successive rows of cells, shown in Figure 1C, is given by

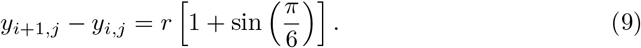

**Fig 1.**
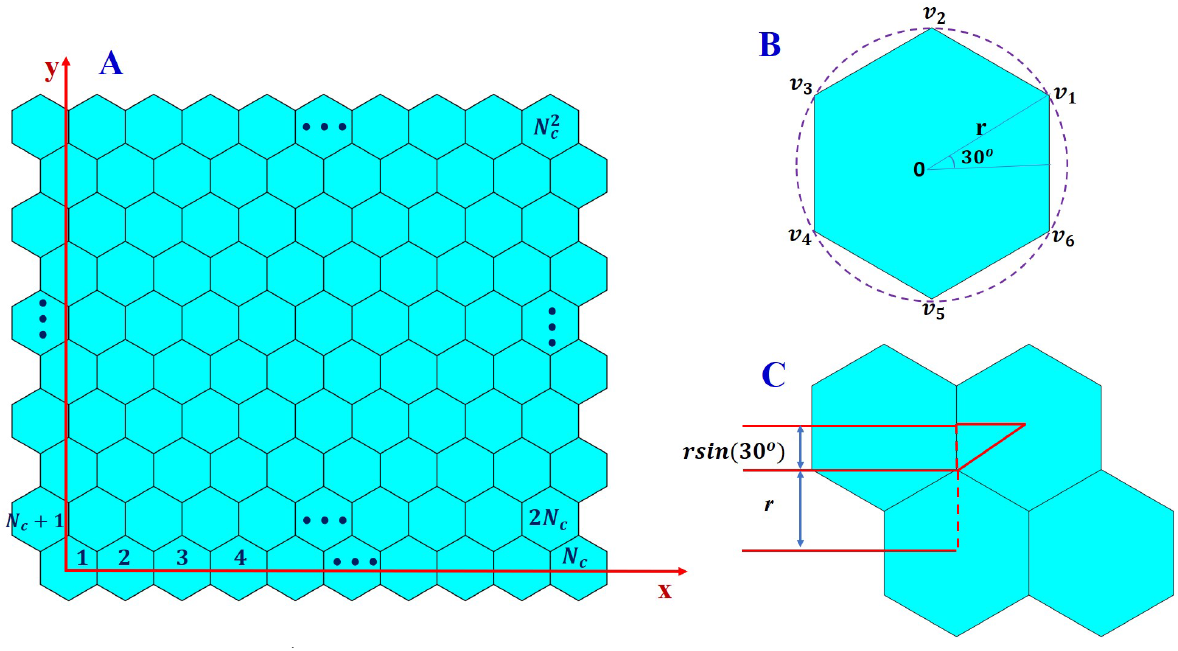
2D tissue: Panel A illustrates a two-dimensional tissue composed of hexagonally packed cells and shows the indexing scheme used to uniquely identify each cell. Panel B depicts a single hexagonal cell, indicating the labeling and ordering of its vertices. Panel C illustrates two adjacent rows of cells and shows how the coordinates of the cell centers are determined.

Similarly, the horizontal distance between the centers of two adjacent cells in the same row is

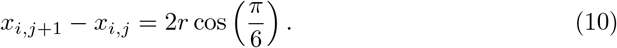

The vertices of a reference hexagonal cell centered at the origin are given by

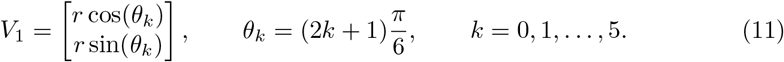

The vertices of any other cell *p* with center (*x*^*p*^, *y*^*p*^) are obtained by translation:

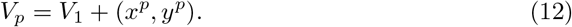

For consistency, the vertices of each cell are numbered counterclockwise, beginning with the vertex located in the first quadrant, as shown in Figure 1B. The vertex and center coordinates are stored for subsequent computations.

#### 2.3.2 Identification of Apical and Filopodia Neighbors

The interaction between cells is obtained by first identifying cells belonging to the apical/junctional and filopodia neighborhoods of any given cell in the tissue. Cell *q* is said to belong to the apical neighborhood of cell *p* if the two cells are in direct contact and the distance between their centers is less than twice the circumradius *r*, that is

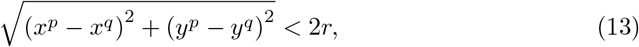

where (*x*^*p*^, *y*^*p*^) and (*x*^*q*^, *y*^*q*^) denote the centers of cells *p* and *q*, respectively.

Assuming that the maximum length any filopodium in the tissue can reach is *D* cell diameters, cell *q* can potentially interact with cell *p* via filopodia if the distance between their centers satisfies

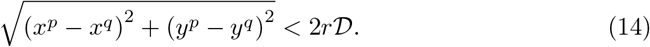

In this case, cell *q* is said to belong to the filopodia neighborhood of cell *p*.

#### 2.3.3 Determination of Filopodia Vectors

The interaction of any pair of filopodia from cells in the same filopodia neighborhood is determined by their orientation and length. In our simulation, we considered two cases: the case where all the filopodia in the tissue are oriented toward a fixed point and the case where filopodia orientation angles are sampled from a normal distribution and scaled to lie within the interval 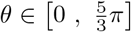 radians, representing an exterior angle of a regular hexagon.

To orient all the filopodia toward a common fixed point, the filopodia angle is defined below and illustrated in Fig. 2.

**Fig 2.**
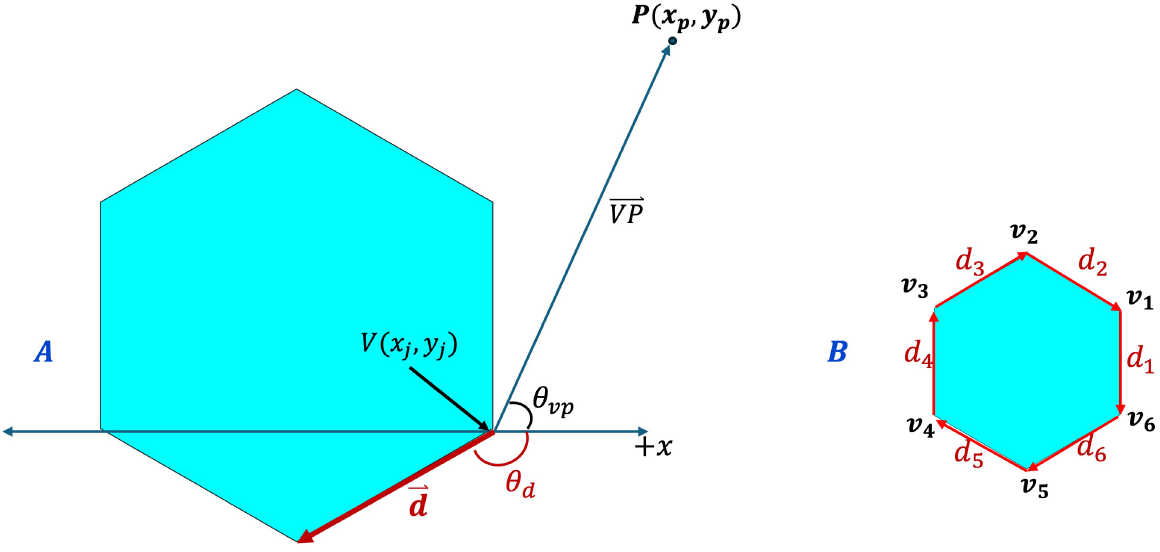
Orientation Angle. (A) Illustration of a filopodium protruding from a cell vertex toward a fixed point. (B) Labeling order of the vertices and edges of a representative cell in the tissue.

##### Definition 2.1

**(Filopodia Angle)** *Let P* (*x*_*p*_, *y*_*p*_) *be a fixed point in the tissue, and let V* (*x*_*i*_, *y*_*i*_) *denote the i*^*th*^ *vertex of a cell. The filopodia angle associated with a filopodium protruding from vertex V and oriented toward the fixed point P is defined as the counterclockwise angle through which the edge vector* **d** *must be rotated to align with the vector connecting V to P*.

Figure 2A illustrates a filopodium protruding from a cell vertex toward a fixed point and the angles formed by the edge vector and the filopodium with the positive *x*-axis, while Figure 2B shows the labeling order of the vertices and edges for a representative cell in the tissue. For the case where the orientation of the filopodia is chosen at random, the particular angle of each filopodium is chosen randomly.

Such precise construction of filopodia has several benefits. First, we are able to investigate the influence of filopodia orientation on patterning. Secondly, we can estimate which pair of filopodia from neighboring cells interact and can therefore facilitate ND signaling.

To compute the orientation angle associated with a filopodium protruding from any vertex of a cell and directed toward a specified fixed point *P* in the tissue, we follow the steps below:

1. Specify the fixed point *P* (*x*_*p*_, *y*_*p*_), identify the vertex *V* (*x*_*i*_, *y*_*i*_) from which the filopodium protrudes, and determine the edge vector **d** originating from the vertex *V* (*x*_*i*_, *y*_*i*_), as shown in Figure 2A.
2. Compute the vector connecting the vertex *V* to the fixed point *P* :

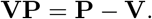
3. Compute the direction angles *θ*_*vp*_ and *θ*_*d*_ that the vectors **VP** and **d** make with the positive *x*-axis, respectively, as shown in Figure 2A. These angles can be computed in MATLAB or Python using the built-in function atan2(y,x):

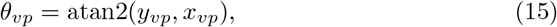

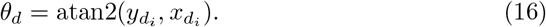
4. Compute the angular difference

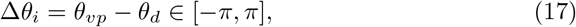

and transform Δ*θ*_*i*_ into the corresponding counterclockwise angle using the modulo operation:

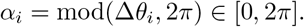

Using the orientation angles and the distribution of filopodia length obtained by solving Equation (4), we compute the direction vector for each filopodium in the tissue, as well as the coordinates at the tips of each filopodium. The direction vectors are used to determine when a pair of filopodia physically intersect, while the coordinates at the tips of the filopodia are used in constructing the network of filopodia during the simulation, as discussed below.

To determine the direction vector of a filopodium, we first compute the unit vector along the corresponding cell edge and then rotate it through the filopodia angle *θ* using the rotation matrix

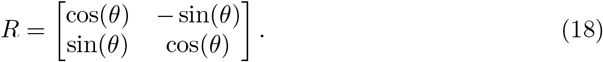

The direction vector of a filopodium is then given by

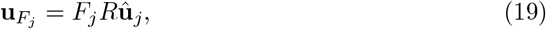

where *F*_*j*_ denotes the length of the *j*^*th*^ filopodium from a given cell and **û**_*j*_ is the unit vector along the cell edge given by

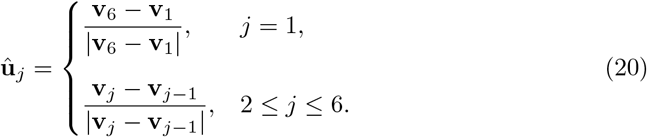

The coordinates of the tip of the *j*^th^ filopodium are given by

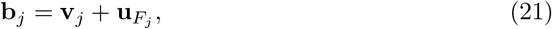

where **v**_*j*_ denotes the coordinates of the *j*^th^ vertex (the base of the *j*^th^ filopodium) of the cell. The numbering of the cell vertices is important, as it determines the ordering of the filopodia and is used throughout the simulation.

#### 2.3.4 Effective Delta Expressed by Neighboring Cells

Long-range Notch–Delta signaling is characterized by the concentration of Delta along each filopodium and the likelihood of Notch activation, which is assumed to be equal for all interacting filopodia. The basal concentration of Delta accounts for about a quarter of the total Delta expressed by each cell, with half available in apical regions and the remaining quarter in the cortex region.

Three-dimensional experimental imaging of fruit flies used in this study revealed that Delta proteins are distributed non-uniformly within cells, with the majority localized at the apical cortex, a smaller fraction at the lateral cortex, and the remainder at the basal cortex [57]. Although these distributions have not been explicitly quantified in the literature, we assume in our simulations that approximately 50% of the Delta within a cell is available for apical interactions, while 25% is available for filopodia-mediated interactions. The remaining 25% is assumed to be localized to the lateral cortex and therefore does not directly participate in signaling interactions. Furthermore, we assume that not every cell-cell interaction results in Notch activation. We therefore define *α*_*a*_ and *α*_*b*_ as the probabilities that apical and filopodia-mediated interactions, respectively, lead to Notch activation. Under these assumptions, the effective Delta contributions from junctional and filopodia neighbors presented in Equation (3) are given by

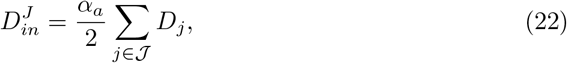

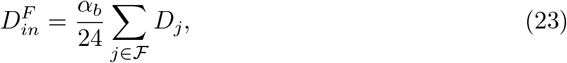

where *J* and ℱ denote the sets of apical and filopodia neighbors of a given cell, respectively.

To compute 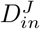 and 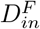 for a given cell, we loop over its apical and filopodia neighbors. The scaling factor *α*_*a*_*/*2 in the computation of 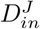 reflects our assumption that only approximately 50% of the Delta expressed by a neighboring cell is available for apical interactions. Similarly, the scaling factor *α*_*b*_*/*24 in the computation of 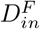 accounts for the assumption that only 25% of a cell’s Delta is available for filopodia-mediated signaling and that this fraction is distributed equally among the six filopodia of the neighboring cell, as illustrated in Figure 3C.

**Fig 3.**
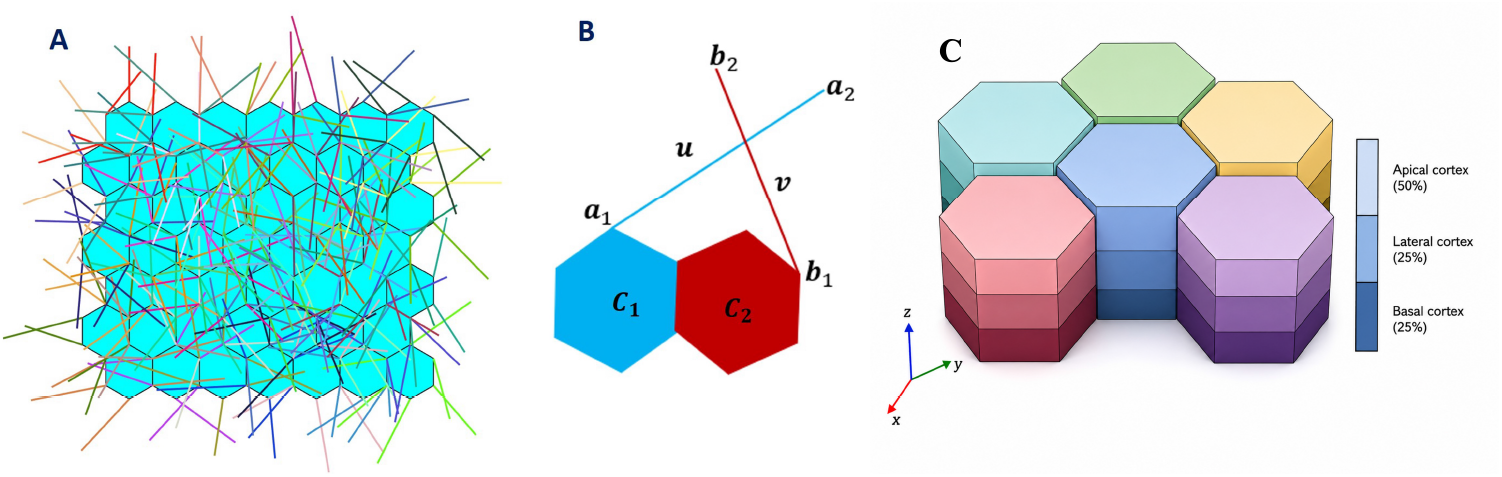
Filopodia Interaction and distribution of Delta: (A) Illustrate a network of filopodia. (B) shows two filopodia from different cells intersecting physically. (C) illustrates the assumed distribution of Delta within a cell, with approximately 50%, 25%, and 25% localized to the apical, basal, and lateral cortices, respectively.

To define the criteria under which any pair of filopodia intersect, we make the following assumptions:

1. All cells are assumed to be hexagonal in shape and identical in size.
2. For each cell, filopodia protrude from all six vertices.
3. Since Notch activation occurs only between two distinct cells, we assume that two filopodia originating from the same cell cannot interact. Therefore, even if they intersect, no Notch activation occurs.
4. A filopodium from a given cell cannot interact with multiple filopodia from the same neighboring cell. For any given filopodium, multiple interactions are only allowed when the interacting filopodia originate from different neighboring cells.
5. Since the protein Ena/VASP, which regulates actin filament dynamics, primarily enhances filament elongation and promotes the formation of unbranched actin structures [36–38], we assume that filopodia protrude along straight lines.

Figure 3A shows a simulated network of basal filopodia for a 7 × 7 grid of hexagonal cells. For any pair of filopodia originating from two distinct cells in the tissue, let **a**_1_ and **b**_1_ be the coordinates at the bases of the two filopodia, and **u** and **v** the direction vectors along the filopodia, obtained by specifying the angles of the filopodia as discussed in Section 2.3.3. As shown in Figure 3B for illustration, the pair of filopodia emanating from the interacting cells *C*_1_ and *C*_2_ lie along the straight lines

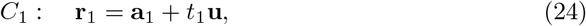

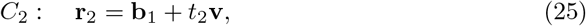

for some constants *t*_1_, *t*_2_ *∈* ℝ. If the two filopodia physically intersect, then there exists a common point of intersection **r**_0_ satisfying

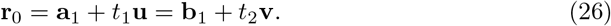

Expanding and simplifying Equation (26) yields the matrix equation

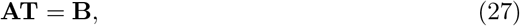

where

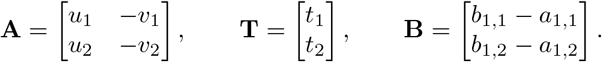

For two filopodia to physically intersect, the matrix **A** in Equation (27) must be non-singular, and the resulting solution must satisfy

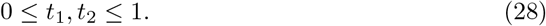

Thus, by defining the system in Equation (27) for any pair of filopodia, one can solve for the constants *t*_1_ and *t*_2_ and determine whether the filopodia intersect. The effective Delta contribution 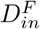 from filopodia interaction is therefore determined by the the filopodia of neighboring cells that make physical contact with the filopodia of a given cell.

#### 2.3.5 Simulating the Notch–Delta Signaling Model

To simulate the Notch–Delta signaling model presented in Equation (1) and Equation (2), we first recognize that the lifetime of the filopodia is much shorter (on average 10 minutes) compared to the time required for the pattern to stabilize (between 16 and 24 hours). Consequently, the filopodia dynamics model must be solved multiple times during the simulation of the Notch–Delta signaling model. We use a smaller time step for the filopodia dynamics model and a larger time step for the Notch–Delta signaling model. The filopodia dynamics model is solved using forward Euler discretization, and filopodia length data are extracted at time points corresponding to the time steps for solving the Notch–Delta model.

At each Notch–Delta time step, the filopodia lengths and orientations are used to compute the effective Delta received by each cell from its neighbors. The effective Delta is then used to update the Notch and Delta concentrations in each cell. When the filopodia lifetime elapses, a new set of filopodia orientations is generated, and the filopodia dynamics model is solved again to obtain updated length data. This process continues until the pattern stabilizes or the prescribed simulation time is reached.

To determine whether the pattern has stabilized, we define a convergence-checking frequency, *T*_*check*_, and a minimum number of unstable cells. We also specify a minimum simulation time before convergence checking begins. At each time step, a moving average is applied to the stored Notch data, which is then used to classify cells as SOP or epithelial cells. The states of the cells are compared between successive iterations, and cells that switch states are recorded. At each convergence check, we determine the number of cells that have switched states at least once during the preceding *T*_*check*_ iterations. If this number falls below the prescribed minimum, the pattern is considered stable, the time iteration loop is terminated, and the final Notch distribution together with the convergence time is reported. Otherwise, the simulation continues until the next convergence check or until the prescribed final simulation time is reached.

The detailed implementation of the coupled Notch–Delta signaling and filopodia dynamics models is summarized in Figure 4. MATLAB-style pseudocode for the simulation steps is provided in the supplementary material.

**Fig 4.**
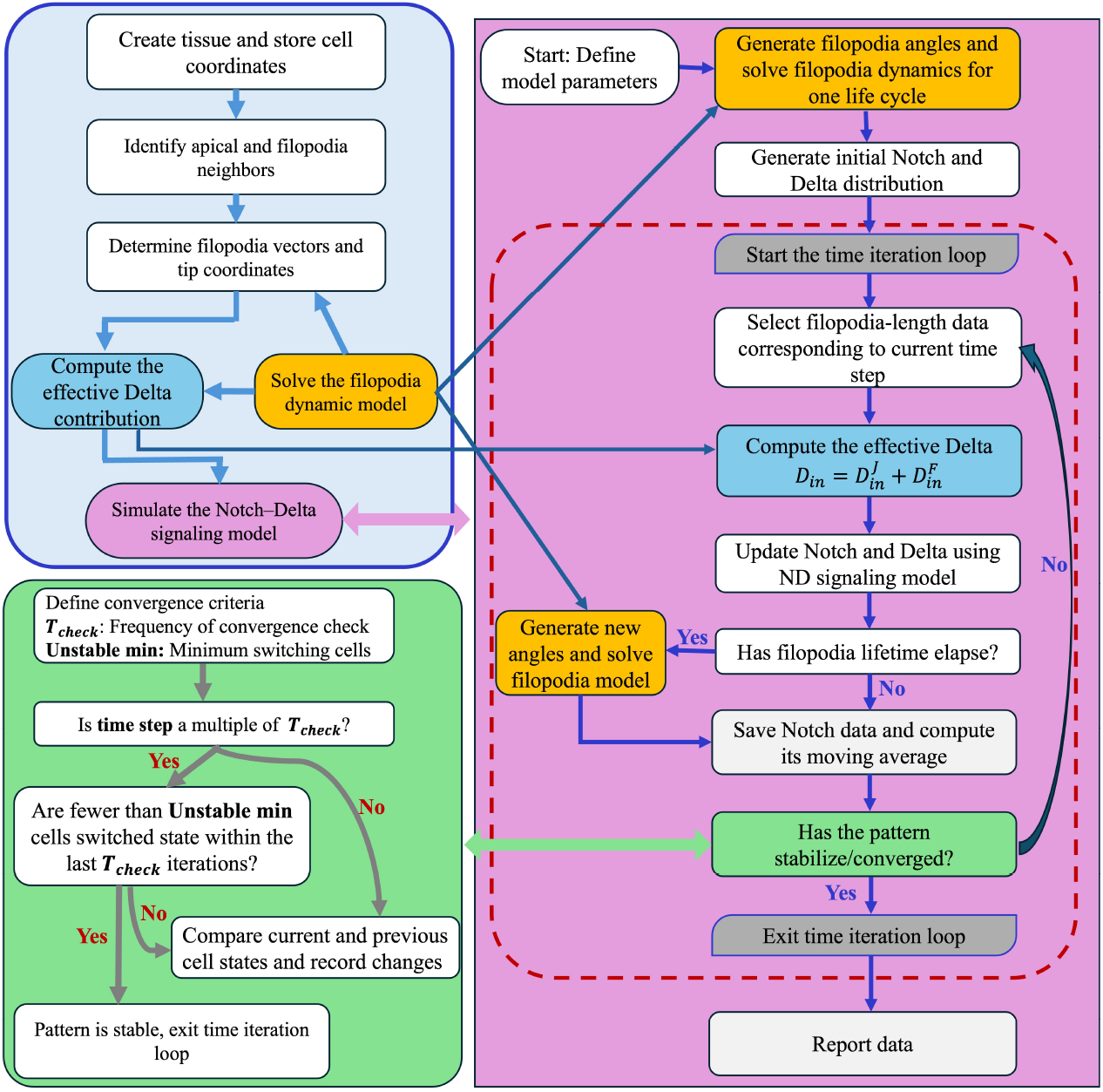
Simulation Flow Chart: The flow chart summarizes the implementation of the Notch–Delta signaling model. The top-left block shows the main components of the simulation, the bottom-left block describes the convergence check, and the right block details the Notch–Delta time iteration loop.

### 2.4 Quantifying SOP Spacing

When the simulation ends, either because a stable pattern was achieved or because the prescribed final simulation time was reached, the resulting Notch distribution is used to quantify the spatial organization of SOP cells using the Relative Sensory Area (RSA) which measures the fraction of the tissue area occupied by SOP cells. It is defined as

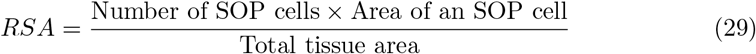

and was introduced in [58]. Although RSA provides information on the overall abundance of SOP cells, it does not account for their local spatial distribution. For example, two tissues may have the same RSA value even if the SOP cells in one tissue are packed into a corner of the tissue, while those in the other are more uniformly distributed.

To address this limitation, we introduce an additional measure called the Local Sensory Area (LSA). The LSA is computed for each SOP cell in the tissue and provides information about the local spatial organization of SOP cells. To compute the LSA, a disc of radius *R* is drawn around each SOP cell, as illustrated in Figure 5. A local fraction of SOP cells within the disc centered at the SOP cell with index *i* is then computed as

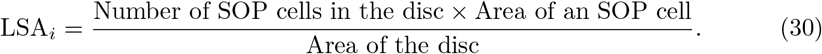

**Fig 5.**
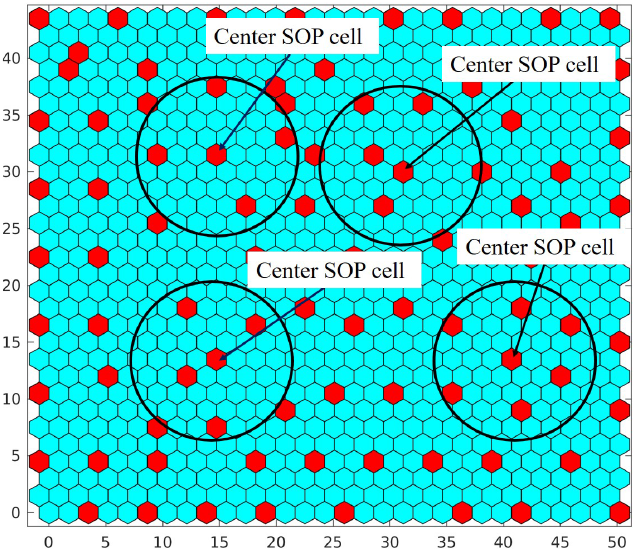
Method of computing LSA: The figure illustrates how the Local Sensory Area (LSA) is computed for each SOP cell in the tissue.

This procedure produces a distribution of LSA values for a given tissue. The mean and standard deviation of the LSA distribution provides additional insight into the spatial arrangement of SOP cells and offers a more robust characterization of tissue organization than RSA alone.

### 2.5 Graphical User Interface

In order to make our algorithm for SOP patterning more accessible to scientists without a background in computer programming, we wrapped it in a Graphical User Interface (GUI), which can be accessed via the web or through a Python package. Figure 6 shows what our GUI looks like when a user initially visits the webpage or runs the package.

**Fig 6.**
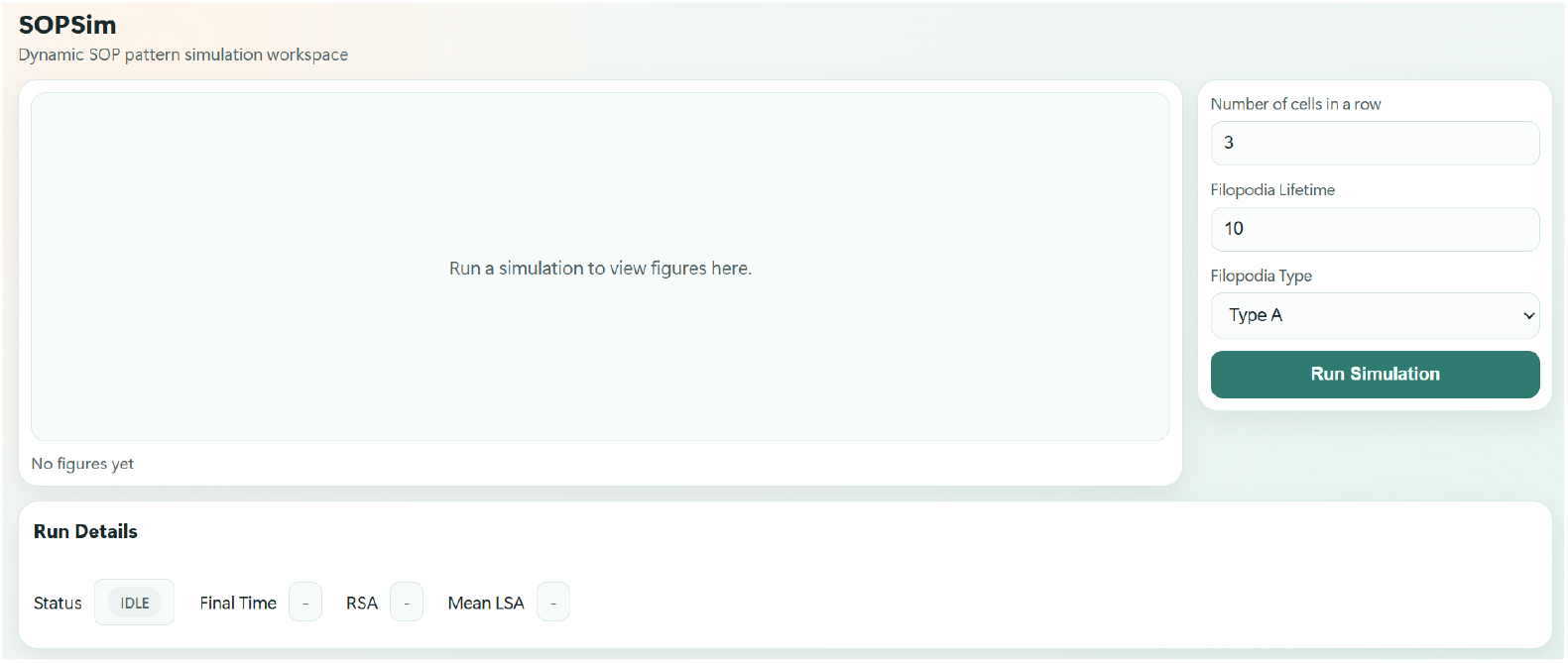
Graphical User Interface: This image showcases a version of the landing page of the Graphical User Interface for simulating Notch Signaling. Users have the option to specify input parameters, run the simulation and observe the outputs of the simulation once it has converged.

**Fig 7.**
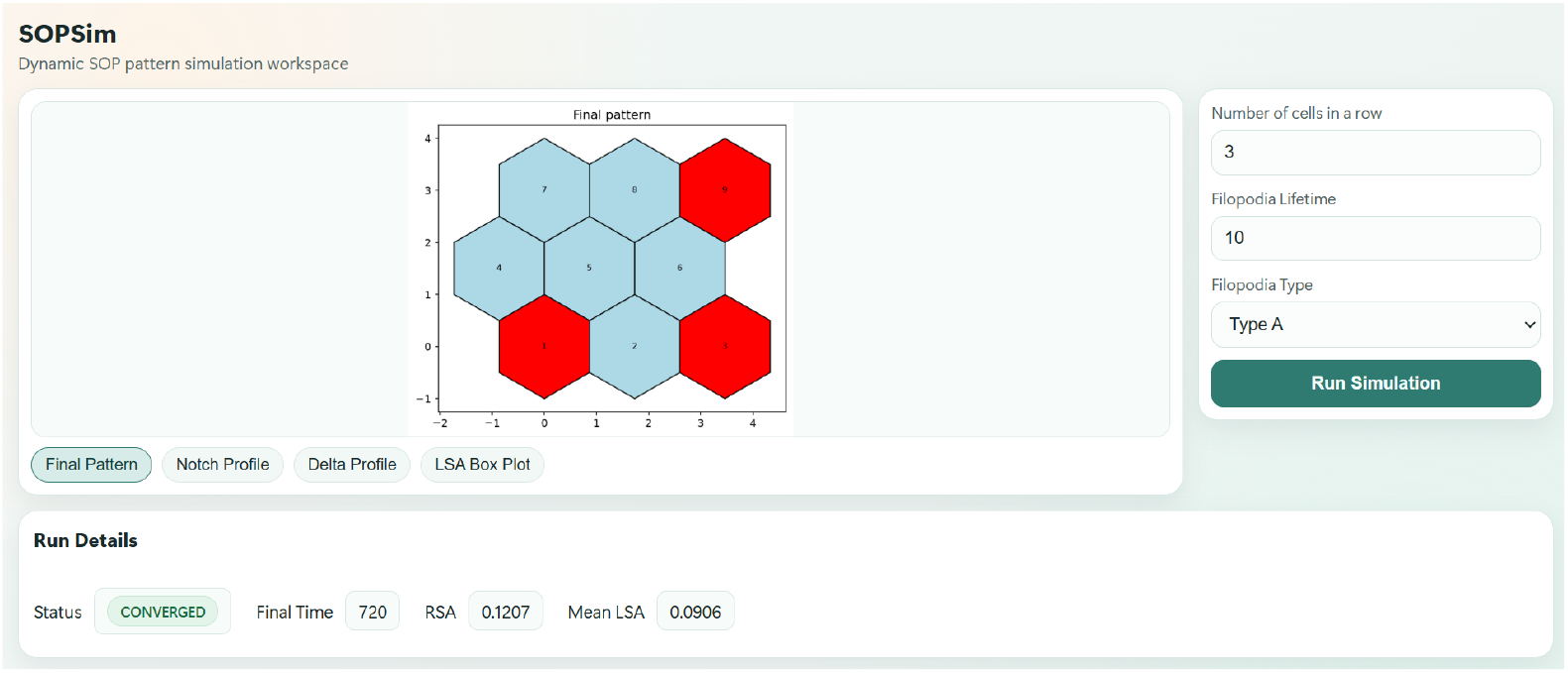
GUI in Converged State: In this figure, we show the user what the GUI looks like after a simulation has been run and the GUI succesfully converged and produced a final SOP pattern for a 3×3 tissue.

In the top right corner, users are able to specify the dimensions of the tissue using the *Number of cells in a row* parameter. If a user enters say 3, then a tissue with a 3 × 3 array of cells is used for the simulation. The other parameter of note is the *Filopodia Lifetime*, which is used to determine how long a Filopodia will protrude, pause, and retract. For simplicity we assume that, a filopodia with a life time of 10 minutes will protrude for 5 minutes (half of its life time) then retract for another 5 minutes. For a life time exceeding ten minutes, the additional time will be spent in a stationary or pausing phase. Consequently, a filopodia with a life time of 17 minutes, will protrude for 5 minutes, remain stationary for 7 minutes, and then retract for 5 minutes. Our GUI allows for the selection of specific filopodia length profiles, as may be determined from future experiments, through the *Filopodia Type* drop down menu.

Once the simulation converges, the user now has the ability to see details about the simulation they just ran. In addition to the final pattern of SOP cells, the GUI presents the *Final Time* at which the pattern converged, as well as metrics describing the spatial organization of the SOP cells such as the RSA, the mean LSA and the distribution of LSA across all SOP cells in the tissue. Other data, such as the average Notch and Delta profile across all SOP and Epithelial cells are also available to the user.

## 3 Results and Discussion

Our proposed computational model for simulating bristle patterning extends Cohen’s model [15] to include deterministic filopodia length dynamics that permit a more explicit representation of the dynamic basal filopodia network regulating long-range cellular interactions and its effect on bristle patterning. In our computational model, filopodia emerge from the vertices of hexagonal cells, packed within a square tissue, at predetermined angles. They are only permitted to perform lateral movements which regulates their length, with an extension and retraction rate governed by a deterministic model for filopodia length dynamics. Random perturbations in the dynamics of each filopodia allow for heterogeneous lengths across each cell and tissue. We have derived a system of linear equations that can be solved to determine when two filopodia from neighboring cells interact. In this way, we can determine that two cells engage in long range ND signaling if any of their filopodia interact. More specially, we are able to quantify the extent of long range signaling that can take place between distant cells through the number of filopodia contacts they make. This additional granularity in filopodia interactions provides biologically relevant heterogeneity in the amount of ND signaling that can occur between two cells reachable by filopodia, and is different from existing models that assume that any two cells that make filopodia contacts will have the same level of long range ND signaling. Details on the formulation of linear equations for determining when filopodia interact, assigning an orientation for each filopodia, the model for filopodia dynamics and the geometric construction of our hexagonal cell-based tissue are described in the Methods and Materials Section.

Our model for filopodia dynamics is modified from Alex Mogilner’s model [50] to allow for a stationary and retracting phase. We achieved this by scaling the actin polymerization rate by a constant *c* during the stationary phase and an exponentially decreasing factor with decay rate *m*, during the retracting phase. We also included a term for retrograde flow velocity (*V*_*r*_), driven by myosin II contraction and depending on myosin pulling force *F*_*m*_ and drag coefficient, *τ*. An experimental study by Hunter *et al*. [31] reported a filopodia retraction rate of 34.2 ± 22.2 nm*/*s. Although the values of myosin pulling force and the drag coefficient have not been explicitly reported in the literature, their values were chosen such that the resulting retrograde flow rate falls within the experimentally observed range of 34.2 ± 22.2 nm*/*s, during retraction.

Parameters for the scaling factor applied to the actin polymerization rate during stationary and retracting phases were selected to reproduce biologically realistic filopodia length distributions. All parameters appearing in Mogilner’s model were used as reported in [50] and presented in Table 2.

With the exception of the Hill coefficient for Delta-induced activation of Notch, *k* and Notch-dependent inhibition of Delta *h*, all other parameters in the ND signaling model presented in Equation (1)–(3) were obtained from previous work by Cohen et al [15] (see Table 1 for details). Initial simulations carried out with the Hill coefficients *h, k* provided in [15] did not yield reported outcomes, hence we decided to estimate them with our new computational framework. A critical transition parameter in SOP patterning is the Notch concentration threshold at which bipotential cells adopt either an epithelial or SOP cell fate. This threshold influences the timescale of the emergence and convergence of the pattern. Recent models by Cohen et al [15], Hadjivasiliou et al [48] and Reppas et al [49] used different Notch thresholds for cell fate determination, without clear support for their use from experimental data. Also, the SOP pattern was considered to have converged when less than 1% of cells changed state, without explicit experimental support. We begin by determining more data-driven thresholds for cell fate and pattern convergence as well as estimates for hill coefficients *k, h*. Following this, we investigate how filopodia lifetime, length, and orientation influence the spatial organization of SOP cells.

**Table 1.** The table presents parameter values used in simulating the Notch-Delta interaction model defined in Equation (1)–(3).

| **Parameter** | **Value** | **Description** |
|---|---|---|
| *R*<sub>*N*</sub> | 0.1 min<sup>−1</sup> | Rate of Notch production [15] |
| *R*<sub>*D*</sub> | 0.1 min<sup>−1</sup> | Rate of Delta production [15] |
| *μ* | 0.002 min<sup>−1</sup> | Decay rate of Notch [15] |
| *ρ* | 0.002 min<sup>−1</sup> | Decay rate of Delta [15] |
| *a* | 0.01 | Half-saturation constant for Notch activation by Delta [15] |
| *b* | 100 | Inhibition strength of Delta production by Notch [15] |
| *k* | 9 | Hill coefficient for Delta-induced activation of Notch (Estimated) |
| *h* | 9 | Hill coefficient for Notch-dependent inhibition of Delta (Estimated) |
| *r* | 1 cell radius | Average cell radius |
| *N*<sub>*μ*</sub> | 1 | Mean Notch level [15] |
| *D*<sub>*μ*</sub> | 1 | Mean Delta level [15] |
| *N*<sub>*σ*</sub> | 0.01 | Standard deviation of Notch levels [15] |
| *D*<sub>*σ*</sub> | 0.01 | Standard deviation of Delta levels [15] |
| *α*<sub>*a*</sub> | 0.1 | Apical weight of expressed Delta [15] |
| *α*<sub>*b*</sub> | 0.1 | Basal weight of expressed Delta [15] |

**Table 2.**
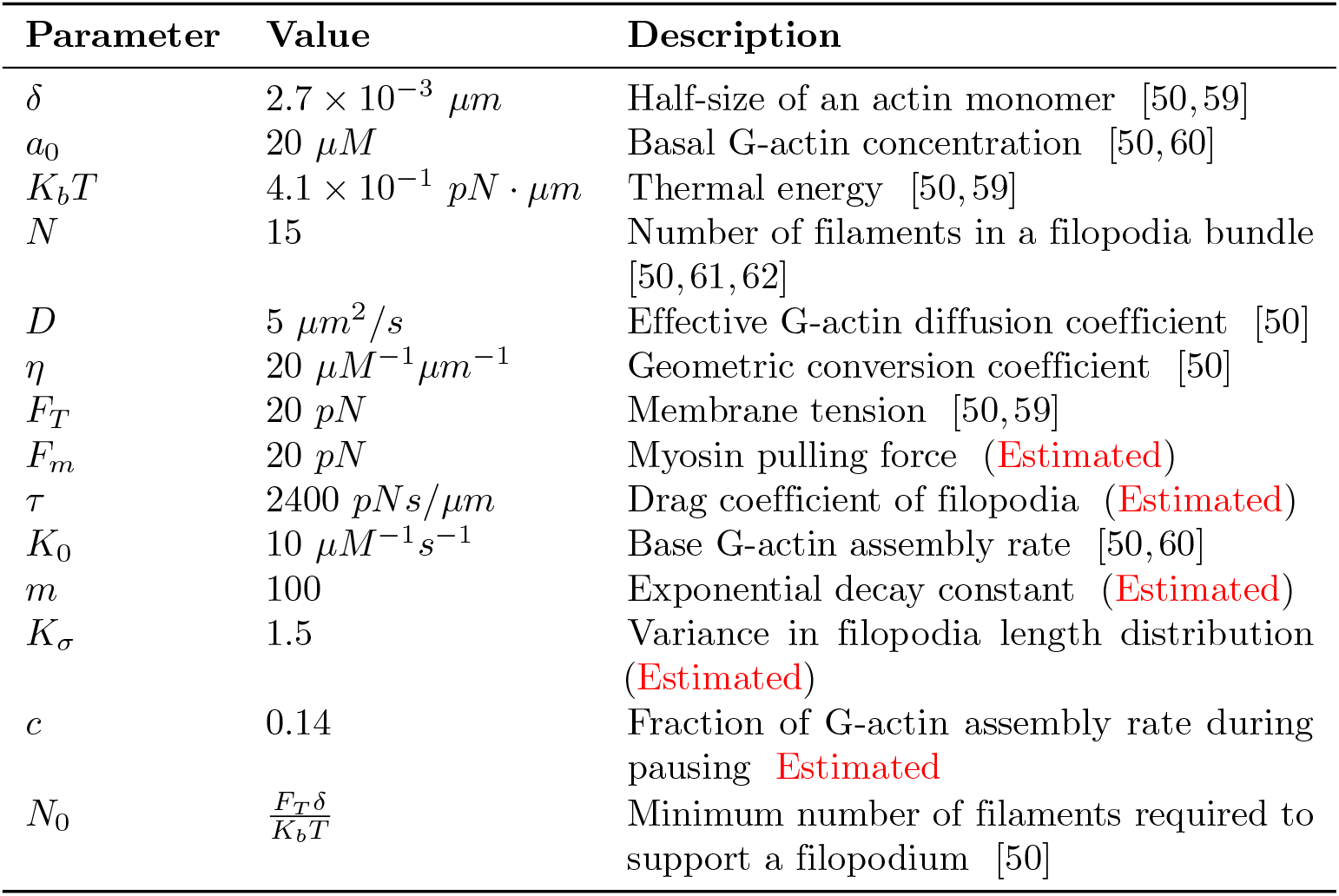
The table presents parameter values used in simulating the filopodia dynamics model described in Section 2.2.

| Parameter | Value | Description |
|---|---|---|
| δ | 2.7 × 10<sup>−3</sup> μm | Half-size of an actin monomer [50, 59] |
| a<sub>0</sub> | 20 μM | Basal G-actin concentration [50, 60] |
| K<sub>b</sub>T | 4.1 × 10<sup>−1</sup> pN · μm | Thermal energy [50, 59] |
| N | 15 | Number of filaments in a filopodia bundle [50, 61, 62] |
| D | 5 μm<sup>2</sup>/s | Effective G-actin diffusion coefficient [50] |
| η | 20 μM<sup>−1</sup>μm<sup>−1</sup> | Geometric conversion coefficient [50] |
| F<sub>T</sub> | 20 pN | Membrane tension [50, 59] |
| F<sub>m</sub> | 20 pN | Myosin pulling force (Estimated) |
| τ | 2400 pNs/μm | Drag coefficient of filopodia (Estimated) |
| K<sub>0</sub> | 10 μM<sup>−1</sup>s<sup>−1</sup> | Base G-actin assembly rate [50, 60] |
| m | 100 | Exponential decay constant (Estimated) |
| K<sub>σ</sub> | 1.5 | Variance in filopodia length distribution (Estimated) |
| c | 0.14 | Fraction of G-actin assembly rate during pausing Estimated |
| N<sub>0</sub> | $\frac{F_T\delta}{K_bT}$ | Minimum number of filaments required to support a filopodium [50] |

### 3.1 Experimental data suggests optimal Notch expression threshold for cell fate determination

To establish a criterion for which cells can be made to adopt either an SOP or epithelial cell fate during simulation, we analyzed data on expression of a Notch transcriptional reporter (NsfGFP) ( [16]) from epithelial (6 cells, 1 tissue), SOP (7 cells, 2 tissues) and switching (35 cells, 3 tissues) cells. Initially, switching cells transiently express an SOP cell fate marker but later express high levels of Notch reporter and adopt an epithelial fate. Cells were imaged every 10 minutes and data collected until nuclear envelope breakdown at the onset of cell division, which coincides pattern stabilization ( [16]). Since the time to cell division varies among cells and the imaging times were not synchronized, times were shifted so that *t* = 0 corresponds to the time at which cell division occurred. This transformation provides a common temporal reference point for the formation of the stable pattern.

Figure 8A shows Notch reporter expression for each cell type, normalized by the maximum expression over the period of observation and averaged across all cells within a given category (i.e. epithelial, SOP, switching). Observe that, during the final 400 minutes to cell division, SOP cells monotonically decrease their Notch reporter expression while epithelial and switching cells increase their Notch reporter expression. We exploited this difference in Notch reporter expression to obtain a threshold that can be used during our computer simulation to distinguish cells moving towards an epithelial fate from those approaching an SOP cell fate. The normalized Notch reporter expression is however not very useful computationally since the maximum expression can only be obtained after the pattern converges. We circumvent this by calculating the moving average in Notch reporter expression for each cell over the period of observation. Specifically, the Notch reporter expression at each time point is scaled by the mean of all preceding values. The mean expression across all cells is shown in Figure 8B. Observe that, within the last 400 minutes to cell division, the average Notch reporter level for SOP cells decreases below one, whereas the average Notch reporter level for epithelial and switching cells remains above one. Based on this observation we determined that, towards the end of patterning, cells with a moving average Notch reporter expression less than 1 will adopt an SOP fate whereas those with a Notch reporter expression greater than 1 will adopt an epithelial cell fate.

**Fig 8.**
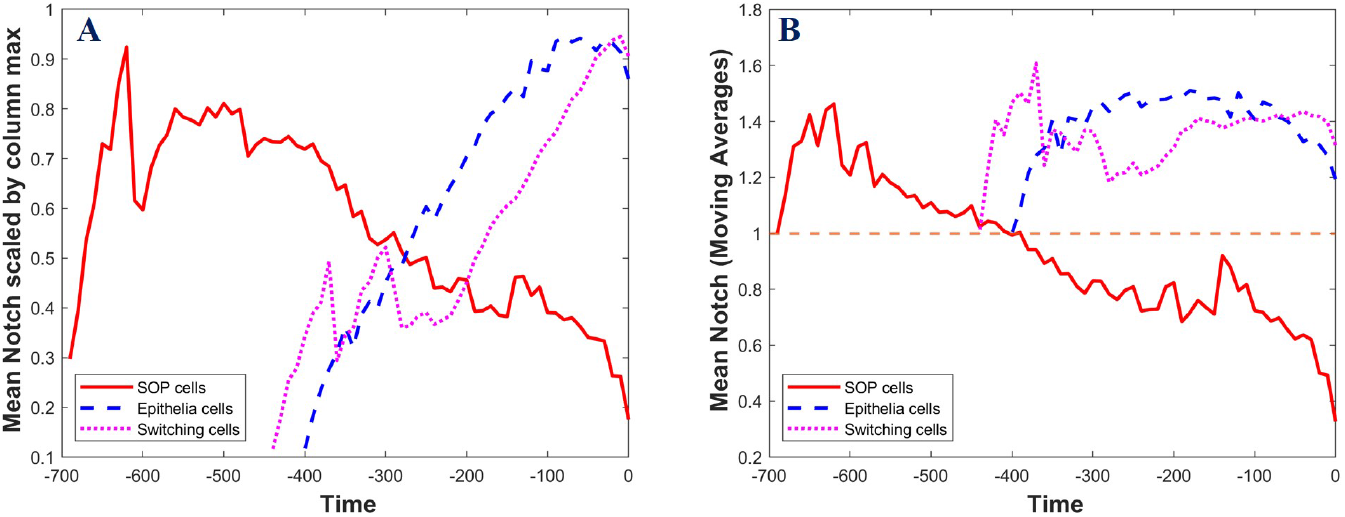
Experimental Notch data: The figure shows trajectory of Notch reporter expression for epithelia, SOP and switching cells up to the time of cell division. In (A), the Notch reporter expression for each cell is scaled by its maximum expression and averaged across all cell in its category. (B) shows the moving average of Notch reporter expression for data presented in A.

### 3.2 Model simulation identifies optimal cell fraction threshold for convergence

A reliable criterion for determining when pattern formation has reached a stable configuration is essential for terminating the simulation efficiently and analyzing how model parameters influence the developmental timescale. We consider the pattern to have converged if fewer than a prescribed percentage of cells, *Cell*_*frac*_ switch state at least once within a fixed observation window of length *T*_check_. Prior to identifying an optimal convergence criteria, we wanted to determine whether our computational model inherently converged. Consequently, we simulated patterning for 3000 minutes, which is well beyond the known experimental window for SOP pattering (960-1440 minutes), and tracked the number of cells changing state within successive windows of *T*_*check*_ minutes. Since SOP patterning is inherently stochastic, due in part to an initial random distribution of Notch and Delta, we performed 25 independent simulations and averaged the number of cells changing state across all simulations. Our results are presented in Figure 9A-E for different values of *T*_*check*_, ranging from 5 minutes to 90 minutes. The corresponding results for *T*_check_ = {120, 150, 200} are provided in the supplementary material, together with the distributions of the switching-cell counts obtained from the twenty-five simulation runs.

**Fig 9.**
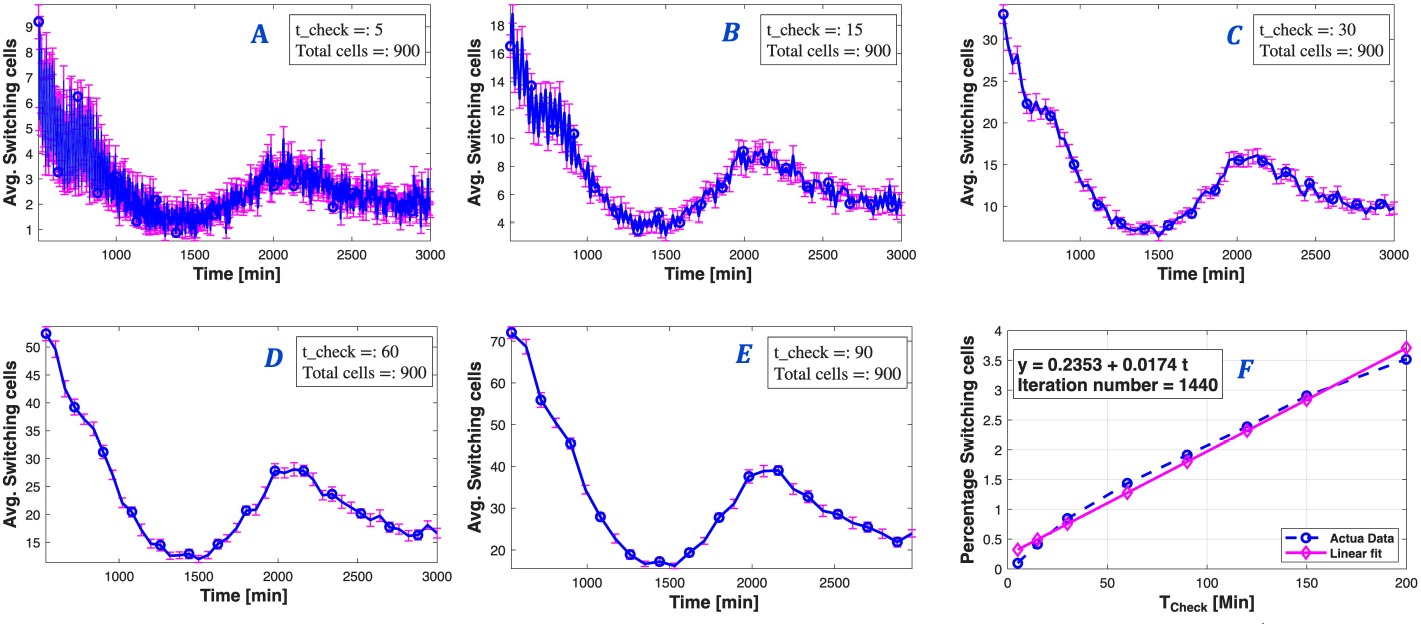
Switching-cell dynamics during pattern formation. Panels A–E show the average number of cells that switch state during each observation window for different convergence-checking intervals, *T*_check_ = {5, 15, 30, 60, 90} iterations. Error bars represent the variability across twenty-five independent simulations. Panel F shows the percentage of switching cells at 24 hours (1440 minutes) of developmental time as a function of *T*_check_, together with the fitted linear trend used to estimate the convergence threshold for different observation frequencies.

Observe that increasing the value of *T*_*check*_ increases the number of cells changing state within that time window. This is expected as a longer time interval will allow for more cells to change state. Remarkably, the number of cells changing state attains a local minimum between 1200 and 1500 minutes, which is consistent with the experimental time scale of SOP patterning. This local minimum can be observed across all values of *T*_*check*_ used. Surprisingly, the number of cells changing state increases after 1500 minutes, reaches a local maximum around 2000 minutes and appears to decrease monotonically towards the 3000 minute mark. We consider the existence of a local minimum within the biological window of stable pattern formation as evidence that the algorithm inherently converges. The progression of patterning dynamics after 1500 minutes suggests that without a biological mechanism, such as cell division, to stop ND-signaling around the 24 hour mark, the pattern may take longer to stabilize.

#### Optimal Hill coefficients and ND signaling sensitivity parameters

The values of Hill coefficients *h, k* supplied by Cohen et al. [15], which govern Notch-dependent inhibition of Delta and Delta-induced activation of Notch, yielded a very dense distribution of SOP cells in our initial simulations. This discrepancy likely arose from the incorporation of explicit filopodia dynamics in the present study. Therefore, we sought to find optimal parameter ranges for (*h, k*) that replicate the known bristle density and time to convergence. We extended this parameter assessment to the apical and basal ND signaling sensitivity parameters, *α*_*a*_ and *α*_*b*_.

Experimental studies suggest that SOP patterns emerge between sixteen and twenty-four hours after pupal formation [29, 63–65]. Bristle density was measured using the relative sensory area (RSA), reported in [58], where the analysis of wild type SOP patterns yielded an average RSA of 0.023. See Figure 11A for the complete distribution of RSA. The (*h, k*) value reported in [15] is (3,3). We expanded this range by varying 0 ≤ *h, k* ≤ 12. Fixing all other model parameters to their base line values reported in the literature and summarized in Table 1 and Table 2, we simulated patterning for various pairs of *h, k* and stored the RSA and convergence time of the final pattern. Figure 10A presents a heatmap of RSA values corresponding to all (*h, k*) pairs sampled. The subset of pairs yielding RSA values below 0.1 and convergence times between 12 and 24 hours are shown in Figure 10B. Observe from Figure 10B that parameter values satisfying *h, k* ≥ 8 produced the most biologically realistic patterns. As a secondary validation, we simulated patterning, first with *h* fixed and *k* varied between 0 and 12 and then with *k* fixed and *h* varied between 0 and 12, storing both average RSA and convergence time. The respective profiles are shown in Figure 10C and D. Observe that the RSA attains a local minimum while maintaining biologically plausible convergence times near (*h, k*) = (9, 9). Consequently, we chose this as our optimal Hill coefficients.

**Fig 10.**
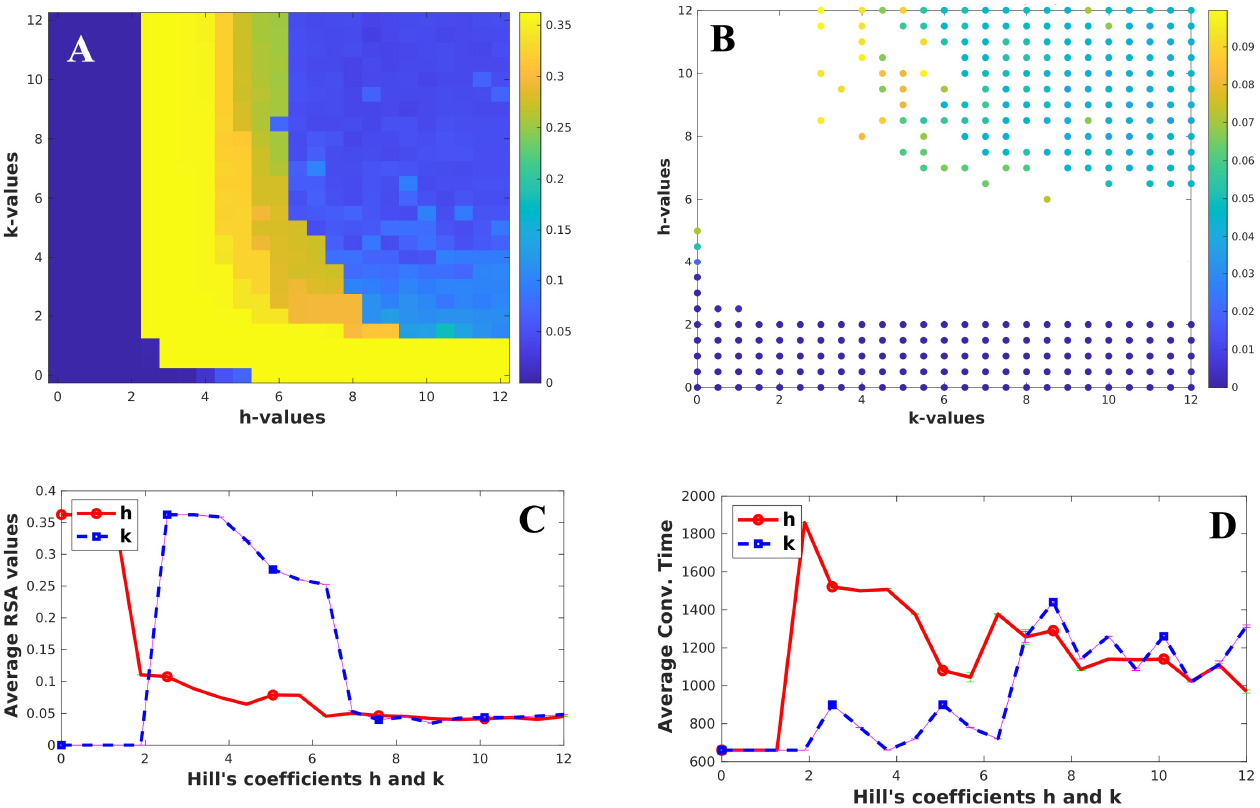
Hill coefficients: The figure illustrates how the Hill coefficient parameters *h* and *k* influence the spatial organization of SOP cells and the convergence time. Panel A presents a heatmap showing the effect of (*h, k*) values on the RSA. Panel B shows the subset of (*h, k*) pairs yielding RSA values less than 0.1 and convergence times between 12 and 24 hours. Panels C and D illustrate the variation of RSA values and convergence times, respectively, as *h* and *k* are varied independently.

**Fig 11.**
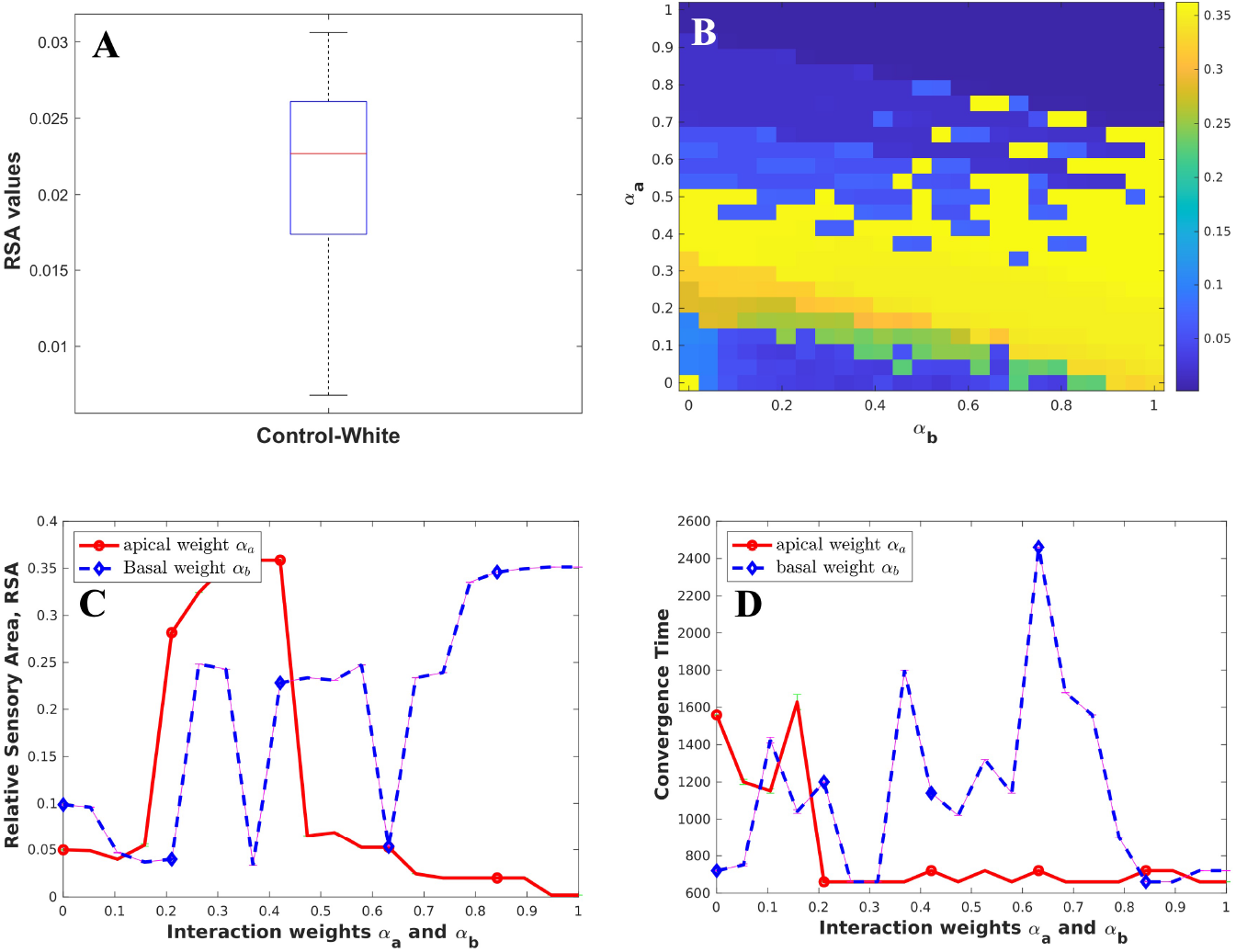
Probability weights: Panel A shows a box plot of RSA values computed from experimental data. Panel B presents a heatmap illustrating how the apical and basal weights, *α*_*a*_ and *α*_*b*_, influence the RSA. Panels C and D show the variation of RSA values and convergence times, respectively, as *α*_*a*_ and *α*_*b*_ are varied independently.

Next, we investigated the effects of the apical and basal weights, *α*_*a*_ and *α*_*b*_, which represent the probabilities that apical and basal interactions lead to Notch activation. The heatmap shown in Figure 11B identifies the range of parameter values that produce RSA values consistent with those observed experimentally. In the study by Cohen et al. [15], the parameter values (*α*_*a*_, *α*_*b*_) = (0.1, 0.1) were used. Our simulations indicate that this choice remains appropriate even after incorporating explicit filopodia dynamics into the model. To further examine the sensitivity of the model to these parameters, we varied *α*_*a*_ and *α*_*b*_ independently within the interval 0 ≤ *α*_*a*_, *α*_*b*_ ≤ 1.0 while keeping the other parameter fixed. Figure 11C and D show the resulting RSA values and convergence times, respectively. The results demonstrate that the RSA is minimized near *α*_*a*_ = *α*_*b*_ = 0.1, with convergence occurring at approximately 1200 minutes (20 hours), in good agreement with the experimentally reported timescale for SOP pattern formation.

### 3.3 Computer simulations replicate experimental evolution of SOP and epithelial Notch expressions

We wanted to assess the ability our simulation framework to replicate the dynamics of Notch expression for SOP and epithelial cells. To achieve this, we first identified the SOP and epithelial cells from the final state of the simulation. For each cell, the Notch level was normalized by its maximum value over time. We then separately computed the average normalized Notch level for the identified SOP and epithelial cell populations.

Figure 12A shows the temporal evolution of the average Notch levels in SOP and epithelial cells. Consistent with the experimental observations shown in Figure 8A, the average Notch level in SOP cells decreases over time, whereas the Notch level in epithelial cells increases. This progressive divergence in Notch expression creates a clear distinction between the two cell fates and reflects the emergence of stable patterning.

**Fig 12.**
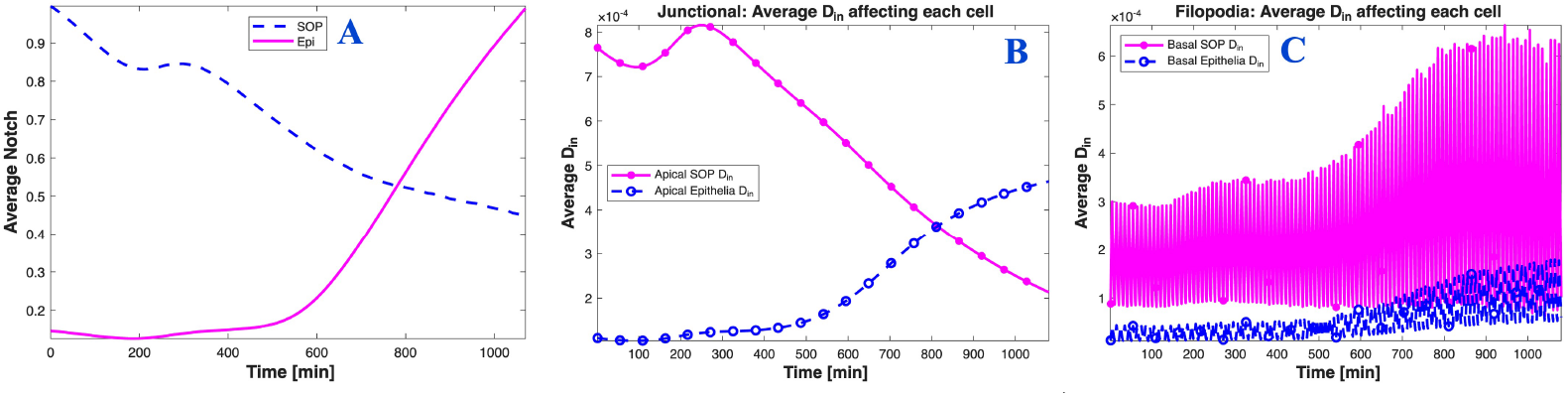
Notch and effective Delta dynamics: Panel A shows the evolution of the average normalized Notch levels in SOP and epithelial cells during the simulation. Panels B and C show the average effective Delta signal, denoted by *D*_*in*_, arising from junctional and filopodial interactions, respectively, acting on SOP and epithelial cells over time.

To further investigate cell fate determination, we examined the evolution of the effective Delta signal, denoted by *D*_*in*_, acting on SOP and epithelial cells. Figure 12B shows the average effective Delta arising from apical/junctional interactions. We observe that the average *D*_*in*_ affecting SOP cells decreases over time, whereas the average *D*_*in*_ affecting epithelial cells increases. These results suggest that SOP cells become surrounded by neighbors expressing lower levels of Delta, while epithelial cells are increasingly influenced by neighbors expressing higher levels of Delta. Such differences in the effective Delta received by the two cell populations likely contribute to the stabilization of cell fates by reducing the likelihood of fate switching once the pattern approaches equilibrium.

The average effective Delta arising from filopodial interactions is shown in Figure 12C. Unlike the junctional contribution, the filopodial contribution exhibits noticeable fluctuations together with a slight increasing trend. On average, the effective Delta received by SOP cells is marginally higher than that received by epithelial cells. The observed fluctuations can be attributed to the highly dynamic nature of filopodia, whose life cycles involve protrusion, retraction, and occasionally pausing. In addition, filopodia continuously change direction over successive life cycles, thereby altering the set of neighboring cells with which they interact. These dynamic behaviors naturally give rise to temporal fluctuations in the effective Delta signal.

To compute the average effective Delta affecting SOP cells (respectively epithelial cells), we summed the effective Delta acting on all SOP cells (respectively epithelial cells) and divided by the corresponding number of cells. Since SOP cells are less abundant than epithelial cells and filopodial neighbors change over time, the average effective Delta for SOP cells is normalized by a smaller population size, resulting in slightly larger values compared to epithelial cells. Overall, these findings provide mechanistic insight into the Notch–Delta signaling dynamics observed *in vivo* when explicit filopodia dynamics are incorporated into the model.

### 3.4 Influence of filopodia length and lifetime on SOP spacing

Experimental studies have shown that filopodia are highly dynamic cellular protrusions, with extension and retraction occurring over timescales of several minutes [15, 46, 66]. Since the average lifetime of filopodia has not been explicitly quantified in the literature, we consider lifetimes ranging from 5 to 20 minutes in order to investigate how filopodia persistence influences the spatial organization of SOP cells. To avoid situations in which long-lived filopodia continue to increase in length indefinitely, we assume in our simulations that filopodia with lifetimes shorter than ten (10) minutes do not undergo a pausing phase. However, if a filopodium persists beyond ten minutes, the additional time beyond ten minutes is assumed to correspond to the pausing phase.

Filopodia are also known to exhibit viscoelastic behavior [50, 51], with several biological and mechanical factors influencing the maximum length they can attain. These factors include membrane tension, myosin contractility, cytoplasmic viscosity, and the concentration of G-actin monomers available for actin filament assembly. In this study, we focus primarily on the role of the basal concentration of G-actin as a factor influencing filopodia length and examine how changes in G-actin availability affect filopodia length and, consequently, SOP spacing.

Figure 13 illustrates the filopodia length profiles, demonstrating how the lifetime and basal G-actin concentration influence the temporal evolution of filopodia length and the maximum length attained. The figure also presents the corresponding average maximal lengths obtained under different filopodia lifetimes and basal G-actin concentrations. We observe that increasing the basal concentration of G-actin leads to longer filopodia. Biologically, filopodia protrusion occurs through the polymerization of G-actin monomers into actin filaments that push the membrane outward. Increasing the availability of G-actin therefore enhances filament assembly, resulting in greater protrusive growth and longer filopodia.

**Fig 13.**
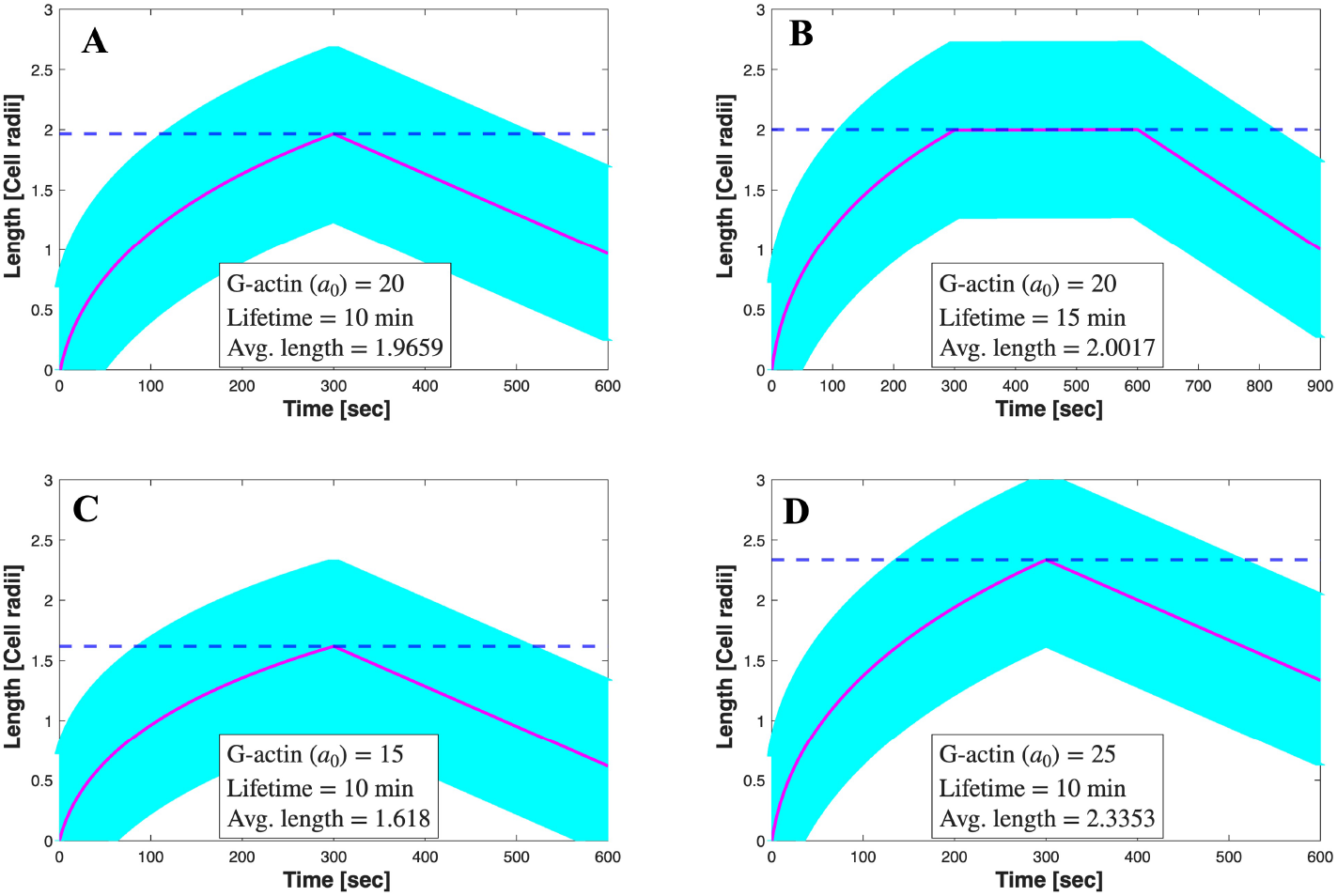
Filopodia length: The figure shows the average maximal length attained by filopodia as the lifetime and basal concentration of G-actin are varied. In panels A and B, the filopodia lifetime was varied to examine the filopodia life cycle and the maximum length attained. In panels C and D, the basal concentration of G-actin was varied to investigate its impact on the maximum length attained by filopodia.

Similarly, increasing the filopodia lifetime results in a longer pausing duration, thereby providing additional time for filopodia to communicate and interact with neighboring cells. In our simulations, however, complete retraction of filopodia was not achieved, which may be attributed to the choice of certain parameter values.

Nevertheless, since the primary role of filopodia is to facilitate communication between non-adjacent cells, the partial retraction observed at the end of each filopodia lifetime may be sufficient to disrupt long-range interactions before the initiation of a new filopodia life cycle. Consequently, this limitation is unlikely to substantially affect the primary objective of the study, namely, investigating how filopodia length and lifetime influence SOP spacing.

Figure 14A and B show, for illustrative purposes, the simulated final SOP–epithelial patterns obtained when all model parameters are fixed at the values listed in Table 2, except for the basal concentration of G-actin regulating filopodia length, which is set to 20 *µM* and 25 *µM*, respectively. The corresponding LSA distributions are shown in Figure 14C and D, together with text boxes displaying the G-actin concentration and filopodia lifetime used as well as the RSA and the average LSA values obtained. We observe that the resulting patterns become progressively finer as the basal concentration of G-actin increases. This suggests that increasing filopodia length enhances long-range interactions, thereby producing a sparser and more organized distribution of SOP cells.

**Fig 14.**
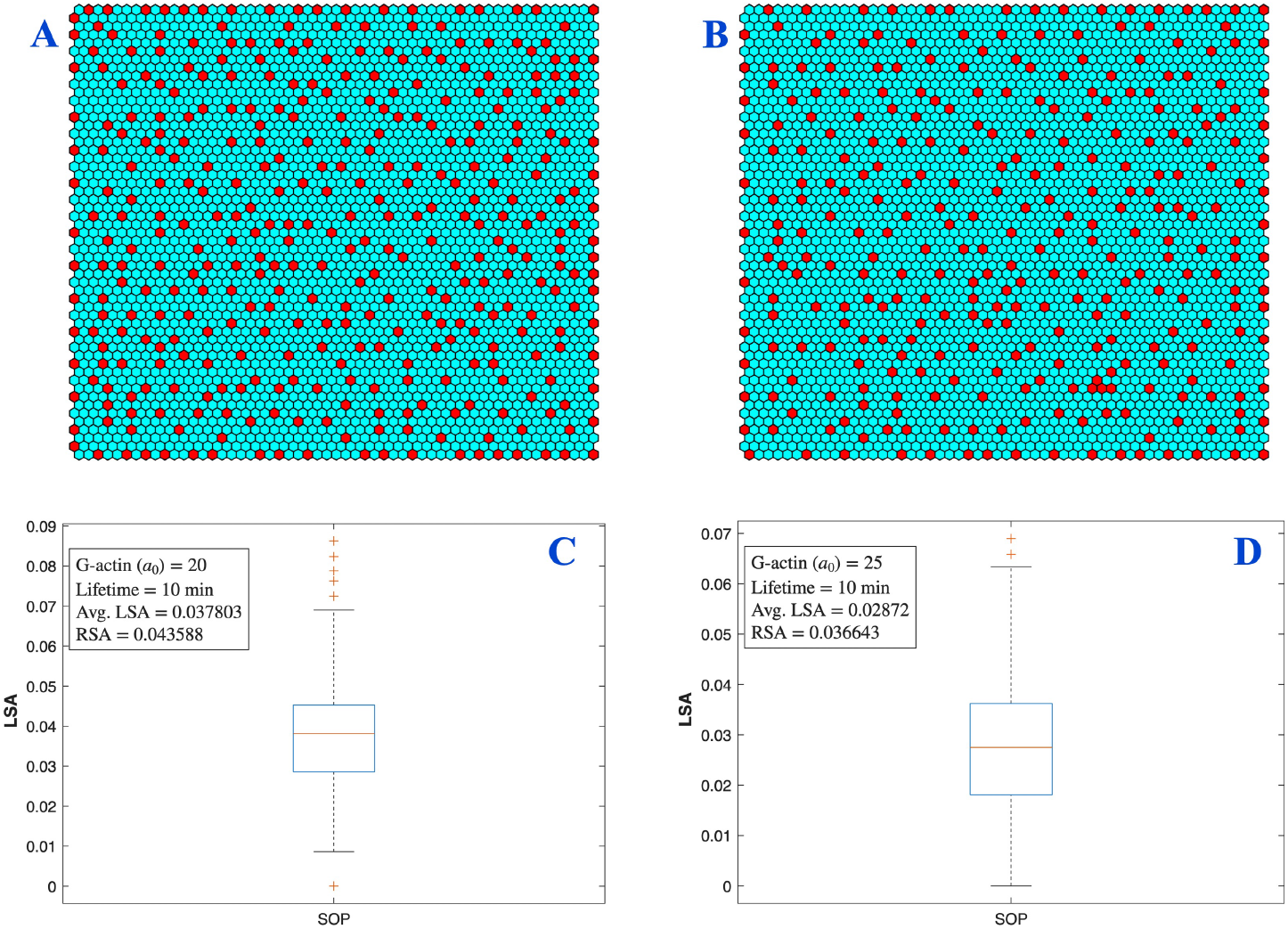
Final pattern: Panels A and B show the simulated final spatial patterns of SOP cells corresponding to basal G-actin concentrations of 20 *µM* and 25 *µM*, respectively. Panels C and D show the corresponding boxplots of the LSA distributions, with the associated convergence times and RSA values displayed in the text boxes.

At the start of each simulation, the initial Notch and Delta levels in the tissue are sampled from normal distributions with prescribed means and standard deviations. To ensure that the observed trends are not artifacts of random initialization, we repeated the simulations twenty-five (24) times for each parameter set and computed the average RSA, LSA, and convergence time.

Figure 15A and B show the average RSA and LSA values, together with error bars representing the standard error across repeated simulations, as the basal concentration of G-actin and the filopodia lifetime are varied, respectively. The results in Figure 15A indicate that increasing the basal concentration of G-actin, which corresponds to longer filopodia, enhances the spatial organization of SOP cells. Specifically, both RSA and LSA decrease as the basal concentration of G-actin increases, suggesting that longer filopodia are able to reach and interact with more distant cells. This increase in long-range lateral inhibition promotes a sparser distribution of SOP cells within the tissue.

**Fig 15.**
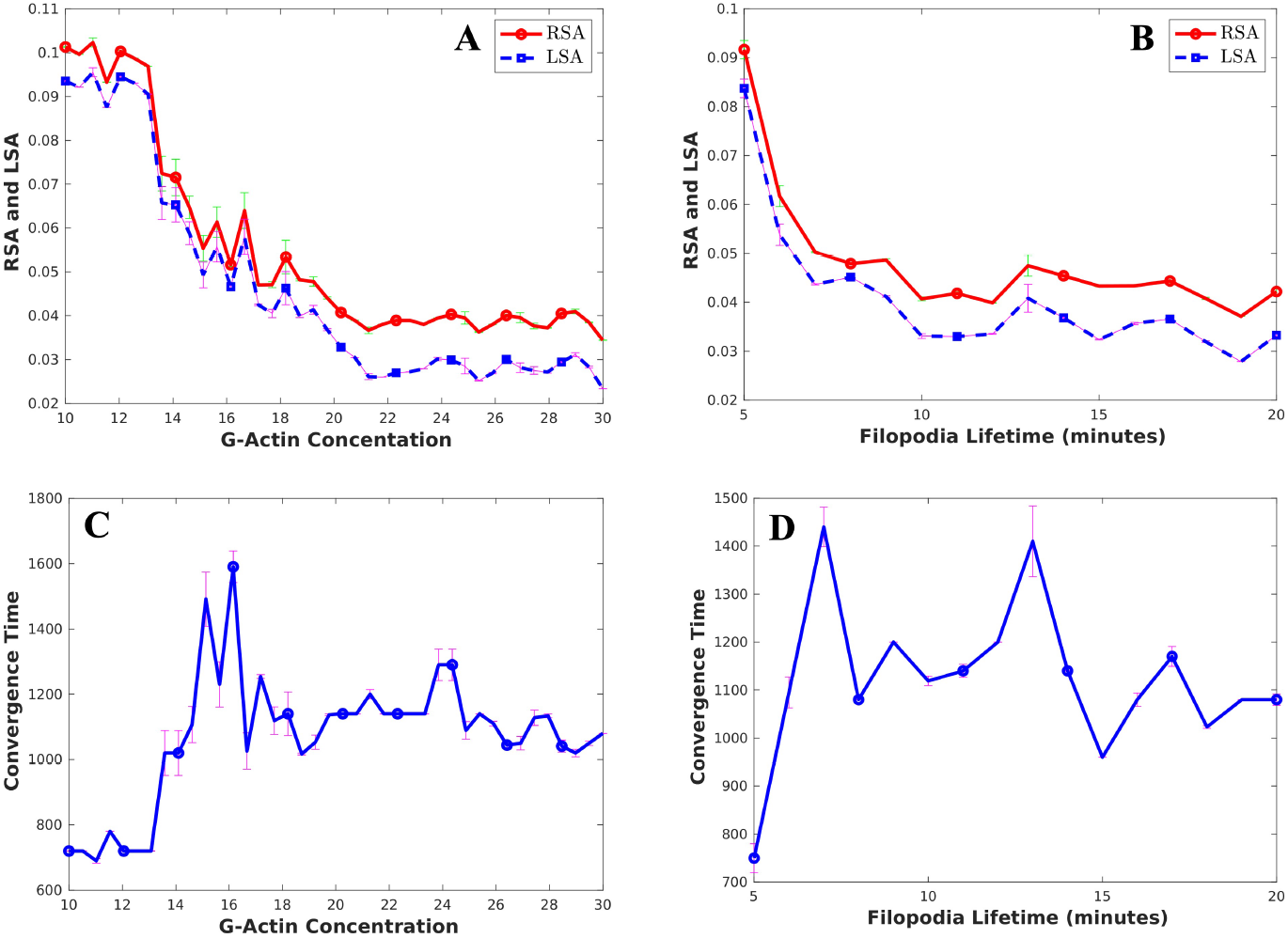
LSA-RSA trend: The figure illustrates the effects of filopodia lifetime and filopodia length on SOP spacing. Panels A and B show the variation of the average RSA and LSA values as the basal concentration of G-actin and the filopodia lifetime are varied, respectively. Panels C and D show the corresponding average convergence times.

The results in Figure 15B show a rapid initial decrease in both RSA and LSA values as the filopodia lifetime increases up to approximately 10 minutes, followed by a more gradual decrease for longer lifetimes. This behavior can be explained by our modeling assumption that filopodia with lifetimes shorter than 10 minutes do not undergo a pausing phase. Consequently, increasing the lifetime within this regime effectively increases the maximum filopodia length, thereby strengthening long-range interactions and promoting a more spatially organized SOP pattern. For lifetimes exceeding 10 minutes, the additional time is allocated to pausing, during which changes in filopodia length are negligible. The continued, but slower, decrease in RSA and LSA values suggests that prolonged filopodia persistence can still enhance signaling interactions; however, its influence is considerably weaker than that of increasing filopodia length. Overall, these results indicate that filopodia length is the primary factor governing SOP spacing, while extended pausing contributes only modestly to pattern refinement.

The corresponding convergence times are shown in Figure 15C and D. In both panels, we observe a sharp increase in convergence time as the basal G-actin concentration and filopodia lifetime increase from small values, followed by relatively constant convergence times, with minor fluctuations, for larger parameter values. Since low G-actin concentrations and short lifetimes both correspond to shorter filopodia, these results suggest that patterns stabilize more rapidly when long-range communication is limited, a result that agrees with the observations of M. Cohen et al. [15] that reducing the lifetime of filopodia resulted in fast pattern formation. As filopodia become longer and are able to establish interactions with more distant cells, additional signaling interactions must be resolved before the system reaches a stable configuration, resulting in longer convergence times. Importantly, the average convergence times for all parameter combinations remain within the experimentally reported time frame for SOP pattern formation, indicating that the spatial patterns generated by the model are biologically realistic.

### 3.5 Effect of filopodia orientation on SOP spacing

To investigate the effect of filopodia orientation on pattern formation, we fixed all model parameters and examined the resulting SOP distribution when all filopodia in the tissue were oriented toward the upper right corner, the upper left corner, or the center of the tissue. These results were compared with the case in which filopodia orientations were assigned randomly. As shown in Figure 16, orienting all filopodia toward a fixed point produces a denser SOP pattern (Figure 16*A*–*C*), whereas randomly oriented filopodia produce a more refined and sparser pattern (Figure 16*D*). To verify that this observation was not specific to the selected directions, we performed additional simulations in which filopodia were oriented toward several other locations within the tissue (results not shown). In all cases, the resulting patterns remained denser than those obtained with randomly oriented filopodia. We also varied the filopodia length and lifetime; however, these changes did not significantly alter the pattern when all filopodia were directed toward a fixed point. The small variations observed in the average LSA and RSA values, for directed filopodia, are attributed to the random initialization of the Notch and Delta concentrations.

**Fig 16.**
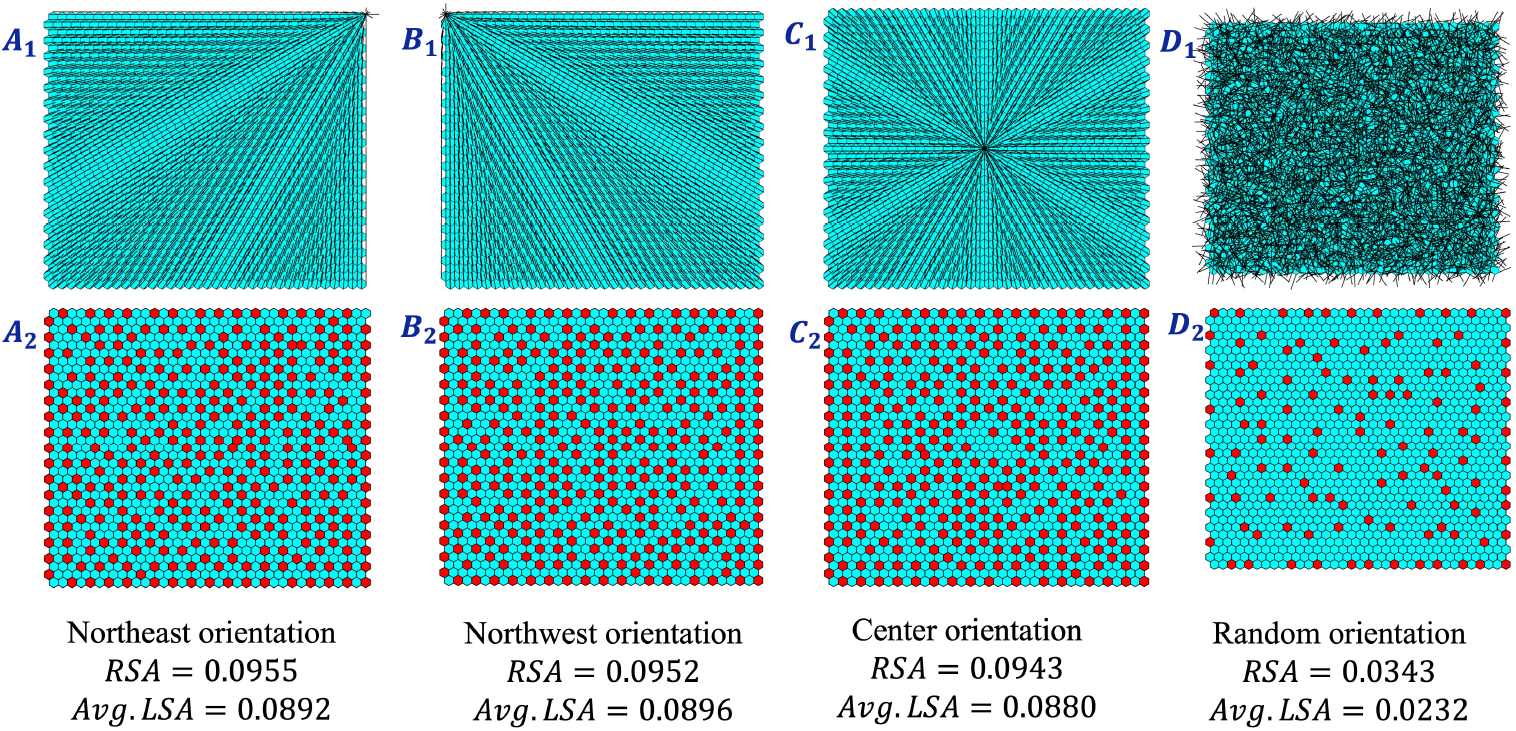
SOP spacing for oriented filopodia: Panels *A*_1_, *B*_1_, and *C*_1_ show the networks of filopodia oriented toward the upper right, upper left, and the center of the tissue, respectively. Panels *A*_2_, *B*_2_, and *C*_2_ show the corresponding SOP patterns. Panels *D*_1_ and *D*_2_ show the network of randomly oriented filopodia and the corresponding SOP pattern. The same parameter values were used in all simulations. The average LSA and RSA values for each simulation are reported in the text boxes below the panels.

To better understand why oriented filopodia produce denser SOP patterns, we monitored the effective Delta (*D*_*in*_) arising from filopodia-mediated interactions throughout the simulation. At each time step, the effective Delta received by every cell through filopodia interactions was stored. After the simulation, these values were separated into SOP and epithelial cell populations, and the mean effective Delta for each population was computed at every time point. Scaling these mean values by the corresponding number of SOP or epithelial cells yields the average effective Delta from filopodia-mediated interactions influencing each SOP or epithelial cell presented in Figure 17. As shown in Figure 17*D*, randomly oriented filopodia generate substantially higher levels of effective Delta than filopodia oriented toward a fixed point, with the average effective Delta being approximately twice as large throughout the simulation. This suggests that orienting filopodia toward a common target reduces the number of long-range contacts formed between cells, thereby weakening long-range Notch–Delta signaling. Consequently, cell-cell communication becomes increasingly dominated by local junctional interactions, leading to the denser SOP patterns observed in Figure 16. The oscillations in the curves arise from the continuous protrusion and retraction of filopodia during the simulation. In Figure 17, the final point on the horizontal axis corresponds to the time at which the pattern reaches convergence. Consistent with the convergence-time results shown in Figure 15*C* and *D*, reducing long-range communication results in faster pattern stabilization and a denser final SOP distribution.

**Fig 17.**
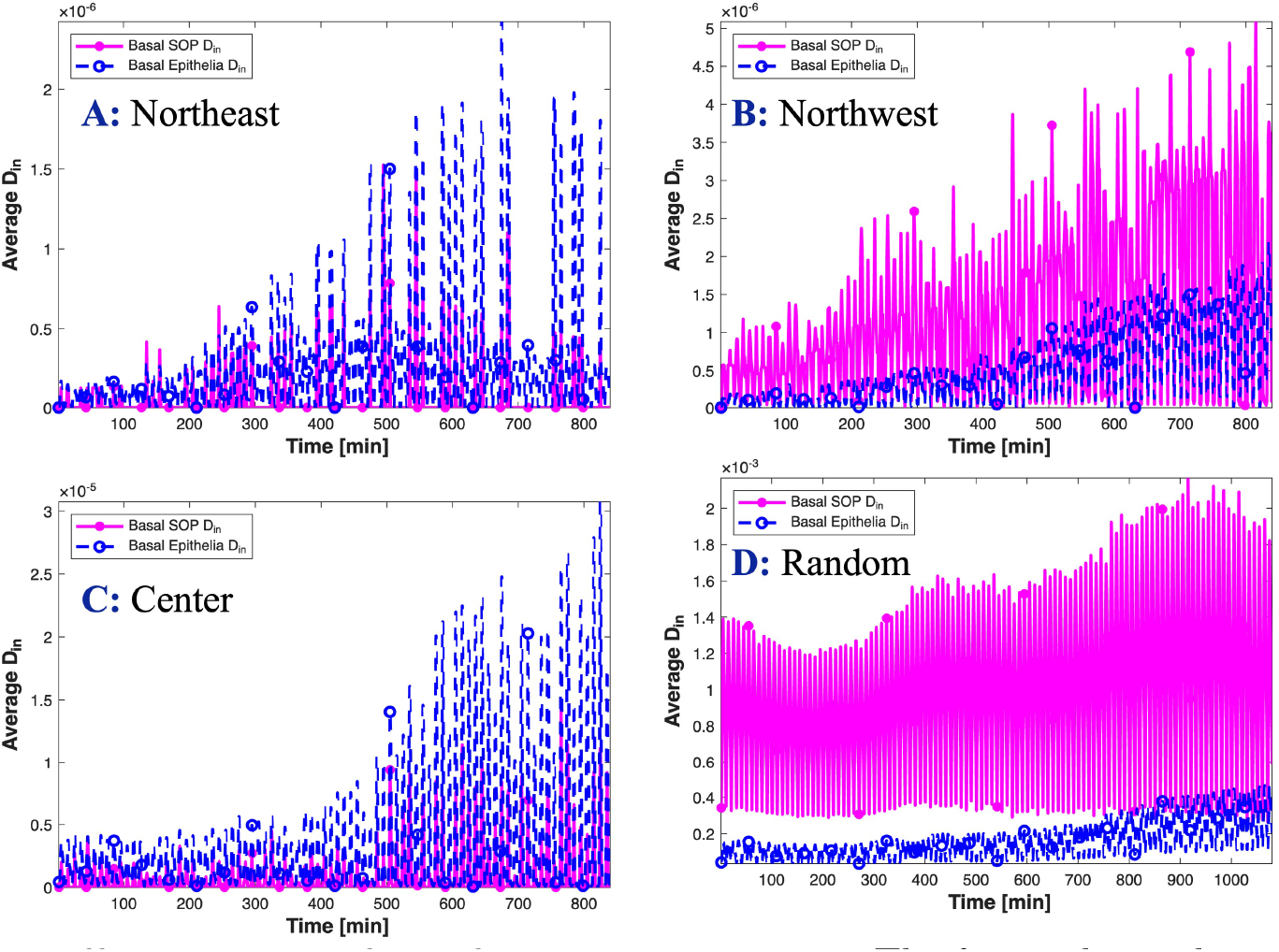
Effective Delta from filopodia neighbors: The figure shows the average effective Delta received through filopodia-mediated interactions by individual SOP and epithelial cells over the course of the simulation. Panels *A, B*, and *C* correspond to filopodia oriented toward the upper right, upper left, and the center of the tissue, respectively, while Panel *D* corresponds to randomly oriented filopodia. In each panel, one curve represents the average effective Delta acting on SOP cells and the other represents the average effective Delta acting on epithelial cells.

## 4 Conclusion

In this study, we extended the Notch–Delta signaling framework of Cohen et al. [15] by incorporating an explicit model for filopodia dynamics that captures the protrusion, pausing, and retraction behaviors observed *in vivo*. This extended framework enabled us to investigate how the dynamic properties of filopodia influence long-range signaling interactions and sensory organ precursor (SOP) pattern formation in developing tissues. In addition, we developed a deterministic criterion for identifying physical intersections between filopodia, thereby providing a geometrically consistent mechanism for long-range signaling interactions.

To quantify spatial organization beyond global pattern measures, we introduced a new metric termed the *Local Sensory Area* (LSA), which characterizes the local distribution of SOP cells within the tissue. We also established a threshold criterion for distinguishing between SOP and epithelial cells using experimental Notch data. The coupled Notch–Delta and filopodia dynamics models were solved numerically using a MATLAB implementation. The MATLAB-type pseudocode is presented in the supplementary material. The actual MATLAB code is publicly available in the GitHub repository https://github.com/williamannan/Notch-Signaling-Simulation/tree/main. A Python implementation of the model with a graphical user interface (GUI) is also available, enabling users without computational expertise to specify a few parameters, simulate the model, and visualize the results.

Our numerical investigations validated the apical and basal interaction weights previously proposed by Cohen et al. [15] and identified biologically realistic values for the Hill coefficients governing Notch activation and Delta inhibition. The simulations demonstrated that filopodia length plays a central role in regulating SOP spacing. In particular, increasing the basal concentration of G-actin resulted in longer filopodia, enhanced long-range interactions, and produced more refined and spatially organized SOP patterns. We further found that increasing filopodia lifetime promotes pattern refinement, although its effect depends on how lifetime influences filopodia dynamics. For lifetimes below approximately 10 minutes, increasing lifetime effectively increases the maximum filopodia length and substantially enhances SOP spacing. For longer lifetimes, the additional time is primarily allocated to a pausing phase during which filopodia length changes very little. In this regime, further increases in lifetime continue to improve spatial organization, but the effect is considerably weaker. Collectively, these results indicate that filopodia length is the dominant factor governing SOP spacing, while prolonged filopodia persistence provides a secondary contribution by modestly strengthening long-range signaling interactions. The investigation of the effect of filopodia orientation on SOP spacing revealed that, orienting all the filopodia in the tissue towards a fixed point reduces the effective long range communication thereby producing denser SOP pattern.

The simulations also revealed an important relationship between filopodia-mediated communication and the timescale of pattern formation. When filopodia are allowed to protrude in random directions, filopodia length plays a crucial role in determining how quickly the SOP pattern stabilizes. Specifically, short filopodia, which have a limited ability to establish long-range contacts, produce patterns that stabilize relatively quickly. In contrast, longer filopodia increase the extent of long-range signaling, leading to more refined SOP patterns but requiring additional time for the tissue to reach a stable configuration. Despite these differences, the convergence times obtained across all parameter regimes remained within the experimentally reported time window for SOP pattern formation, supporting the biological relevance of the model predictions.

Although the present model captures key aspects of filopodia-mediated signaling, several simplifying assumptions remain. In particular, apical and filopodial interactions are represented through probability weights rather than explicit molecular signaling events. In future work, we aim to incorporate the transport and accumulation of the Notch intracellular domain (NICD) at base of filopodia of signal-receiving cells. Since NICD regulates downstream gene expression, including Delta suppression and Notch production, such an approach would provide a more mechanistic representation of Notch activation and may yield deeper insight into the mechanisms governing SOP pattern formation.

## Acknowledgments

GH is supported by NIH R35GM150782. EA and GH are supported by the Joint NSF/NIGMS Mathematical Biology Program, R01GM152810.

